# Stable Cell Clones Harboring Self-Replicating SARS-CoV-2 RNAs for Drug Screen

**DOI:** 10.1101/2021.11.04.467291

**Authors:** Shufeng Liu, Chao-Kai Chou, Wells W. Wu, Binquan Luan, Tony T. Wang

## Abstract

The development of antivirals against severe acute respiratory syndrome coronavirus 2 (SARS-CoV-2) has been hampered by the lack of efficient cell-based replication systems that are amenable to high-throughput screens in biosafety level 2 laboratories. Here we report that stable cell clones harboring autonomously replicating SARS-CoV-2 RNAs without S, M, E genes can be efficiently derived from the baby hamster kidney (BHK-21) cell line when a pair of mutations were introduced into the non-structural protein 1 (Nsp1) of SARS-CoV-2 to ameliorate cellular toxicity associated with virus replication. In a proof-of-concept experiment we screened a 273-compound library using replicon cells and identified three compounds as novel inhibitors of SARS-CoV-2 replication. Altogether, this work establishes a robust, cell-based system for genetic and functional analyses of SARS-CoV-2 replication and for the development of antiviral drugs.

**IMPORTANCE:** SARS-CoV-2 replicon systems that have been reported up to date were unsuccessful in deriving stable cell lines harboring non-cytopathic replicons. The transient expression of viral sgmRNA or a reporter gene makes it impractical for industry-scale screening of large compound libraries using these systems. Here, for the first time, we derived stable cell clones harboring the SARS-CoV-2 replicon. These clones may now be conveniently cultured in a standard BSL-2 laboratory for high throughput screen of compound libraries. This achievement represents a ground-breaking discovery that will greatly accelerate the pace of developing treatments for COVID-19.

## INTRODUCTION

The single-stranded, positive-sense SARS-CoV-2 RNA genome is approximately 30 kb in length and comprises a short 5’ untranslated region (UTR), 13 open reading frames (ORFs), a 3’ UTR, and a poly(A) tail. Through discontinuous transcription events, the virus makes at least nine canonical subgenomic RNAs (sgmRNA), which encode structural and accessory proteins (1). The genomic RNA (gRNA) harbors two large ORFs, ORF1a and ORF1ab, which are initially translated into two polyproteins, pp1a and pp1ab, and subsequently processed by viral proteases to produce 16 non-structural proteins (Nsp) that form the viral replication complex and confer immune evasion (2–4).

Viral replication and translation machinery offer some of the most promising targets for antiviral drug development. For example, the main protease (Nsp 5) and the viral RNA-dependent RNA polymerase (Nsp 12) of SARS-CoV-2 are primary targets for antiviral discovery because they are responsible for cleavage of replicase polyproteins 1a/1ab and for virus replication, respectively. A cell-based system that harbors the minimally essential SARS-CoV-2 replication and translation machinery is highly desirable because it enables simultaneous screening of inhibitors of multiple viral proteins in a biosafety level 2 setting. To this end, subgenomic viral RNA molecules called replicons are often designed to autonomously replicate in cells without generating infectious virus. Current SARS-CoV-2 replicon systems, however, do not permit persistent replication in cell lines due to intrinsic toxicity (3, 5-8). The limited time window allowable for detection of replication, the inability to generate master and working cell banks for lot consistency, and the challenge to scale up for industrial processes, make it impractical to apply transient replicon systems in high-throughput screening (HTS) of large compound libraries. Here, we describe, for the first time, the derivatization and characterization of stable cell clones harboring autonomously replicating SARS-CoV-2 RNAs. We further demonstrated the applicability of cells harboring SARS-CoV-2 replicon RNA in drug screen.

## RESULTS

### Initial design of SARS-CoV-2 replicon

To generate stable cell clones harboring replicating SARS-CoV-2 RNAs, we first constructed a replicon termed SARS-CoV-2-Rep-NanoLuc-Neo, in which the Spike (S) gene was replaced by a nanoluciferase reporter (NanoLuc), and the Envelope (E) and Membrane (M) genes were replaced with the neomycin phosphotransferase gene (Neo or NeoR) (Fig. 1A top panel). Electroporation of this replicon RNA along with *in vitro* transcribed RNA encoding the nucleocapsid protein (NP), which was previously shown to improve launch efficiency (9–11), into Vero E6, Huh7.5.1, A549 and BHK-21 cells resulted in expression of nanoluciferase to varying extents (Fig. S1A-D). However, no viable clones could be recovered after 21 days of selection in G418, suggesting that active replication of the replicon RNA is either unsustainable or cytotoxic. Huh7.5.1 and BHK-21 cells supported higher nanoluciferase expression than Vero E6 and A549, although we cannot rule out that the differences are attributed to different electroporation efficiency of the four cell lines. Because electroporating two different RNA species into the same cell is inefficient, we created a BHK-21 stable cell clone (BHK-21-NP^Dox-ON^) in which NP is expressed in a doxycycline-inducible manner (Fig. S1E). Electroporation of SARS-CoV-2-Rep-NanoLuc-Neo RNA into BHK-21-NP^Dox-ON^ cells resulted in three neomycin-resistant clones out of four million cells. The resulted clones grew very slowly in the presence of 200 μg/mL G418, and no nanoluciferase activity or viral RNA could be detected, indicating the loss of functional replicon RNA. Altogether, persistent replication of SARS-CoV-2-Rep-NanoLuc-Neo could not be achieved in any of the four mammalian cell lines.

**Figure 1.**
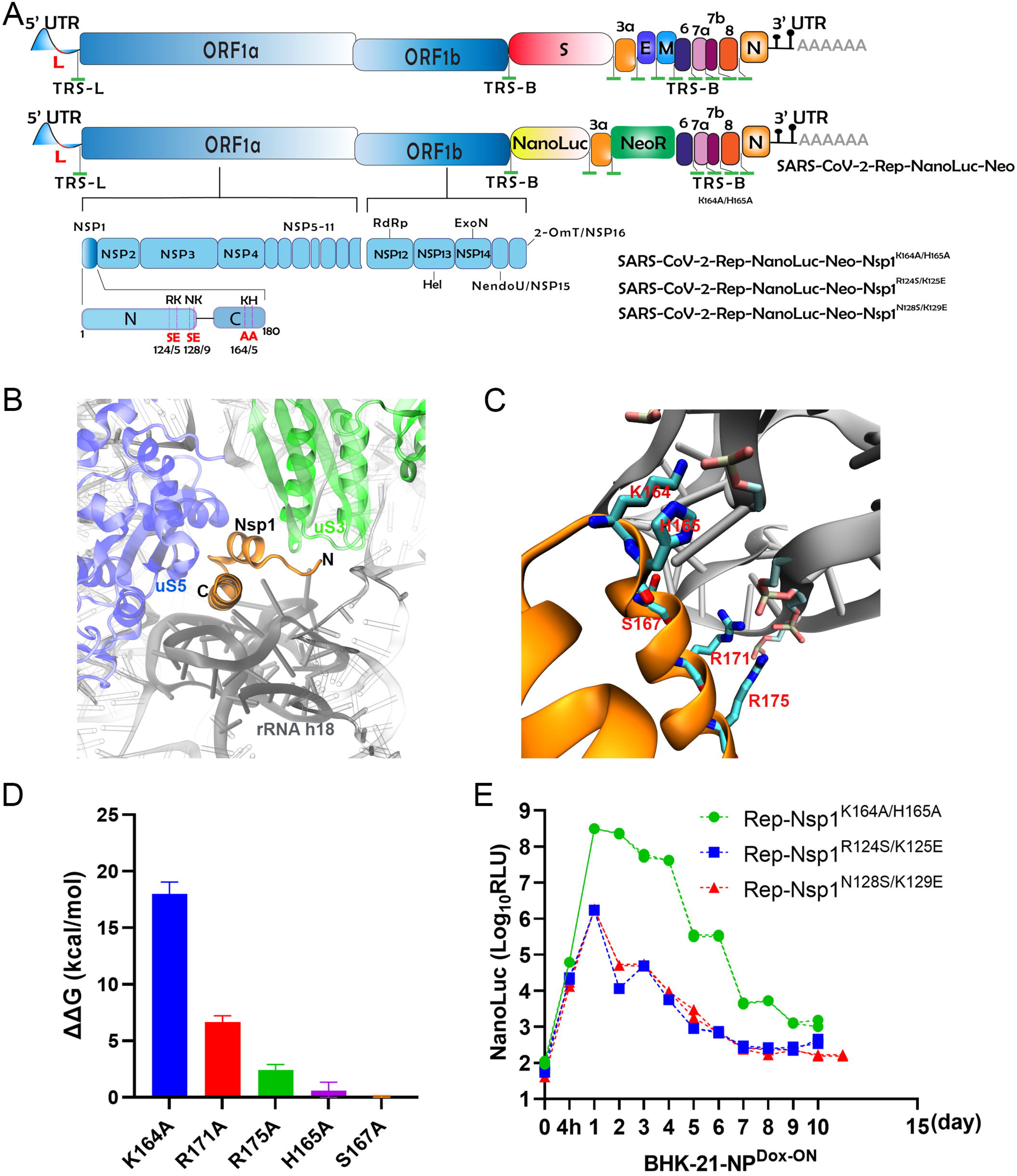
Design and optimization of SARS-CoV-2 replicons. (A) Top, genome organization of SARS-CoV-2. Leader sequence (red), transcriptional regulatory sequence within the leader sequence (TRS-L) and within the body (TRS-B) are highlighted in green. Middle, the design of SARS-CoV-2-Rep-NanoLuc-Neo. Bottom, the Nsp1 mutations were introduced to obtain three more replicons. Figures are not drawn in proportion. (B) Illustration of Nsp1 binding to the small ribosomal subunit (PDB code:7K5I). Nsp1 (orange) binds close to the mRNA entry site and contacts uS3 (green) from the ribosomal 40S head as well as uS5 (blue), and h18 of the 18S rRNA (charcoal gray) of the 40S body. The fragment of rRNA not close to Nsp1 is shown transparently. (C) An enlarged view of the Nsp1 binding area. Critical residues are shown in the stick representation and are highlighted in red. (D) Calculated free energy changes (∆∆G) for various mutations in Nsp1. Positive values indicate unfavorable mutations for the binding between Nsp1 and rRNA. (E) BHK21-NPDox-ON cells were transiently transfected with Rep-NanoLuc-Neo-Nsp1^R124S/K125E^ RNA, Rep-NanoLuc-Neo-Nsp1^N128S/K129E^ RNA and Rep-NanoLuc-Neo-Nsp1^K164A/H165A^ RNA. Nano luciferase was measured at indicated time points post-transfection.

### An improved SARS-CoV-2 replicon

Coronaviruses have evolved a variety of mechanisms to shut off host transcription and translation (12–14). Of all SARS-CoV-2 proteins, Nsp1 causes the most severe viability reduction in cells of human lung origin (15). The carboxyl terminus of Nsp1 folds into two helices, which insert into the mRNA entrance channel on the 40S ribosome subunit, preventing both the host mRNA and viral mRNA from gaining access to ribosomes and consequently shutting down translation (16, 17) (Fig 1.B). The first C-terminal helix (residues 153–160) makes hydrophobic interactions with the 40S ribosomal protein uS5, and interacts with the 40S ribosomal protein uS3 with salt-bridges (e.g. D156-R143 and E159-K148); the second C-terminal helix (residues 166–178) interacts with ribosomal protein eS30 and with the phosphate backbone of h18 of the 18S rRNA via the two conserved arginines R171 and R175 (16). In between the two helices, a conserved KH dipeptide (K164 and H165) forms critical interactions with h18 through H165 stacking between two uridines of 18S rRNA (U607 and U630), and electrostatic interactions between K164 and the phosphate backbone of rRNA bases G625 and U630 (Fig. 1C). Molecular dynamics simulation followed by free energy perturbation calculation predicted that mutations of residues K164, R171, R175, H165, S167 of Nsp1 to alanine will reduce the interaction in the order of impact (Fig. 1D and Fig. S2). We hypothesize that a pair of mutations, such as K164A/H165A that weaken the interaction between C-terminus of Nsp1 and ribosome, will lead to a shorter occupation time of Nsp1 on the ribosome and increase the accessibility of ribosomes to host mRNA. As a result, the Nsp1-mediated toxicity to the host should be alleviated. To test this possibility, we created a new replicon construct SARS-CoV-2-Rep-NanoLuc-Neo-Nsp1^K164A/H165A^ in which K164/H165 were mutated to alanine (Fig. 1A bottom panel). For comparison, we also made two additional replicons, SARS-CoV-2-Rep-NanoLuc-Neo-Nsp1^R124S/K125E^ and SARS-CoV-2-Rep-NanoLuc-Neo-Nsp1^N128S/K129E^, given that both pairs of mutations (R124S/K125E and N128S/K129E) reportedly reduce Nsp1-mediated cell toxicity in a human lung cell line (15). When electroporating into BHK-21-NP^Dox-ON^ cells, all three replicons led to transient expression of nano luciferase (Fig. 1E). However, in the presence of 200 μg/mL G418, only SARS-CoV-2-Rep-NanoLuc-Neo-Nsp1^K164A/H165A^ yielded viable cells (Pool #1), from which 12 stable clones (Clone #2-13) were subsequently derived by limiting dilution. Three independent experiments were performed, and each time only electroporation of SARS-CoV-2-Rep-NanoLuc-Neo-Nsp1^K164A/H165A^ resulted in viable clones in BHK-21-NP^Dox-ON^ cells (Table S1). It is worth mentioning that SARS-CoV-2-Rep-NanoLuc-Neo-Nsp1^K164A/H165A^ also replicated well in Huh7.5.1 cell line although no viable cells could be recovered after G418 selection (Fig. S3). Moreover, we were able to derive at least 4 clones using standard BHK-21 cells albeit at much lower efficiency.

### Characterization of cells harboring SARS-CoV-2-Rep-NanoLuc-Neo-Nsp1^K164A/H165A^

To explore the possibility for SARS-CoV-2-Rep-NanoLuc-Neo-Nsp1^K164A/H165A^ to persist in cells without selection pressure, G418 was subsequently withdrawn from Pool #1 cells after the initial selection. For up to one week there was no significant loss of nanoluciferase expression, a feature that is compatible with drug screen. The level of nanoluciferase decreased by one log after 10 days culturing without G418 and then by another two logs after 21 days (Fig. 2A).

**Figure 2.**
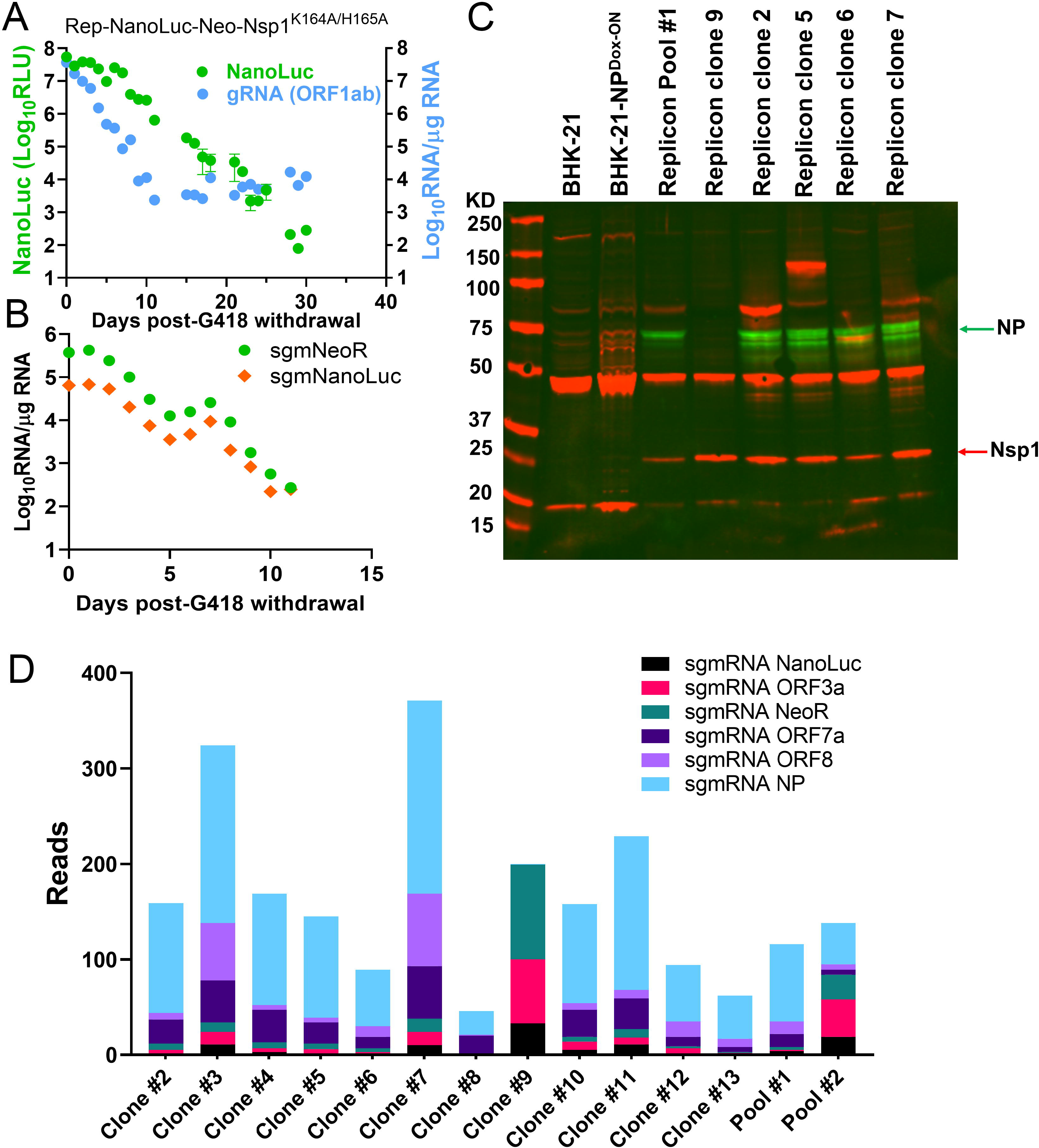
Characterization of replicon cells harboring BHK21-NPDox-ON Rep-NanoLuc-Neo-Nsp1^K164A/H165A.^ (A) Nano luciferase in BHK21-NP Dox-ON replicon cells was measured at given time points following G418 withdrawal. RNA was also extracted at indicate time points and quantified by RT-qPCRs targeting ORF1ab (gRNA in A), or sgmNeoR and sgmNanoLuc (B). (C) Western blot analysis of the SARS-CoV-2 proteins from six representative stable replicon clones. The presence of NP and Nsp1 protein in cell lysates was confirmed. (D) Sequence coverage of canonical sgmRNA species in Pool #1 and Pool #2 replicon cells as well as in each of the 12 stable clones.

Next, we performed quantitative reverse transcription PCR (RT-qPCR) to profile gRNA and sgmRNA species in cells harboring SARS-CoV-2-Rep-NanoLuc-Neo-Nsp1^K164A/H165A^. SARS-CoV-2 transcribes multiple canonical sgmRNAs, including S, E, M, NP, ORF3a, ORF6, ORF7a, ORF7b, ORF8 and ORF10, although multiple studies have found negligible ORF10 expression and very few ORF7b body–leader junctions (1, 18). In replicon cells, S, M, E sgmRNA are replaced with those encoding NanoLuc and NeoR; sgmRNA encoding ORF6 is lost because its transcriptional regulatory sequence body (TRS-B) resides in the M gene, which is also deleted in the replicon RNA. Hence, cells harboring the replicon would express at least six sgmRNAs, namely, Nanoluc, NeoR, ORF3a, ORF7a, ORF8, and NP. We designed primers/probes to specifically amplify the gRNA of the replicon and sgmRNAs of NanoLuc and NeoR. Shown in Fig. 2B and Fig. S4, the temporal expression of gRNA and sgmRNA for the NanoLuc and NeoR genes were clearly observed in Pool #1 cells. Western blotting confirmed the presence of Nsp1 and NP in replicon cell lysates (Fig. 2C and Fig. S4B). To further characterize the replicon gRNA, next-generation sequencing (NGS) was performed on all 12 stable clones. While the replicon gRNA containing Nsp1^K164A/H165A^ was present in all clones, additional synonymous or missense mutations were detected in each clone (Table S2). Among those, Nsp4 R401S substitution was detected in 10 out of 12 clones, and Nsp10 T111I appeared in 6 out of 12 clones. Ongoing research are studying whether if these changes constitute adaptive mutations which had enabled efficient replication in BHK-21 cells. Notably, sequencing the replicon gRNA in clone #9 also revealed a deletion knocking out ORF7a/b, ORF8 and the first 392 amino acids of the NP (Fig. S5), indicating that NP is dispensable for genome replication. The presence of canonical sgmRNA species in each stable clone was also confirmed by NGS, although the method employed in this study could not locate all sgmRNA species due to uneven coverage of sequence reads over different regions (Fig. 2D).

### Application of replicon cells in drug screen

To demonstrate the suitability of SARS-CoV-2-Rep-NanoLuc-Neo-Nsp1^K164A/H165A^ cells in drug screen, we tested a library of 273 compounds for inhibitory effects. These compounds were selected to target Nsp5 (3CLpro), Nsp3 (PLpro), Nsp12 (RdRP), Nsp15, Nsp16, and X domain by virtual screen (Table S3). At 10 μM, nine compounds exhibited more than 50% inhibition based on nanoluciferase expression (Fig. 3A). Three compounds, Darapladib (predicted to target 3CLpro), Genz-123346 (predicted to target Nsp16), JNJ-5207852 (predicted to target Nsp15) (Fig. 3B), were validated in replicon cells and then in three human cell lines A549-hACE2, Calu-3, Caco-2 cells using live virus. Remdesivir (inhibitor of RdRP) and GC376 (3CL protease inhibitor) were included as positive controls. The IC_50_ of these compounds are provided in Figure 3C. Interestingly, all three compounds displayed cell type-specific activity against SARS-CoV-2, which others also noticed with repurposed drugs (19). To further explore the robustness of replicon cells for drug screen, Remdesivir was added to Pool #1 cells on different days following G418 withdrawal. Cells were subsequently incubated for 2-7 days before nanoluciferase was quantified. We observed little difference in terms of the inhibitory strength when Remdesivir was added between 2 and 9 days following G418 withdrawal (Fig. S6). By contrast, the optimal duration of Remdesivir treatment in replicon cells was between 3 and 6 days. Lastly, we sought to evaluate the clonal variability in response to drug treatment. To this end, stable replicon cell clone #3, #5, #7, #9, #11 and #13 were tested for responses to GC376 treatment. Acquired IC_50_ values ranged from 5.9 μM to 13 μM (mean = 9.3 μM, standard deviation = 2.9 μM), suggesting that all clones are potentially suitable for testing drug efficacy (Fig. 4).

**Figure 3.**
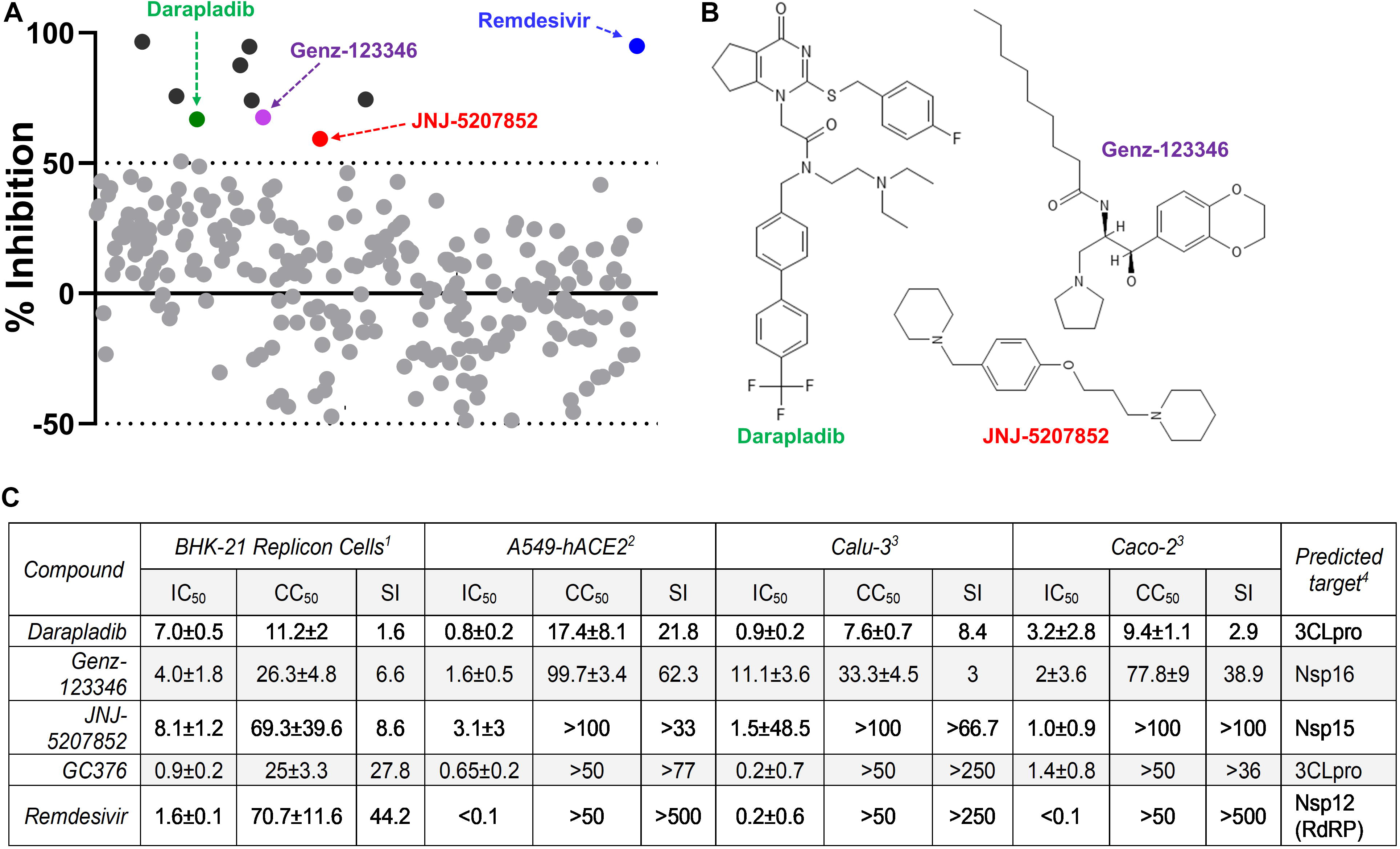
Rep-NanoLuc-Neo-Nsp1K164A/H165A replicon cells for drug screen. A 273-compound library containing virtually identified candidates (see Table S3) was screened in replicon cells (Pool #1) as described in Material and Method. Ten compounds (including Remdesivir) displayed more than 50% inhibition were denoted in black or colored solid circles. (B) Molecular structures of Darapladib, Genz-123346, and JNJ-5207852. (C) Cell type specific activity against SARS-CoV-2 by compounds. ^1^Assays were performed in BHK-21 Pool #1 cells harboring SARS-CoV-2-Rep-NanoLuc-Neo-Nsp1K164A/H165A; ^2^Assays were performed using live SARS-CoV-2 carrying a nanoluciferase and then a firefly luciferase reporter; ^3^Assays were performed using live SARS-CoV-2 carrying a nanoluciferase reporter. All values in μM standard deviation from at least 3 biological or technical replicates.

**Figure 4.**
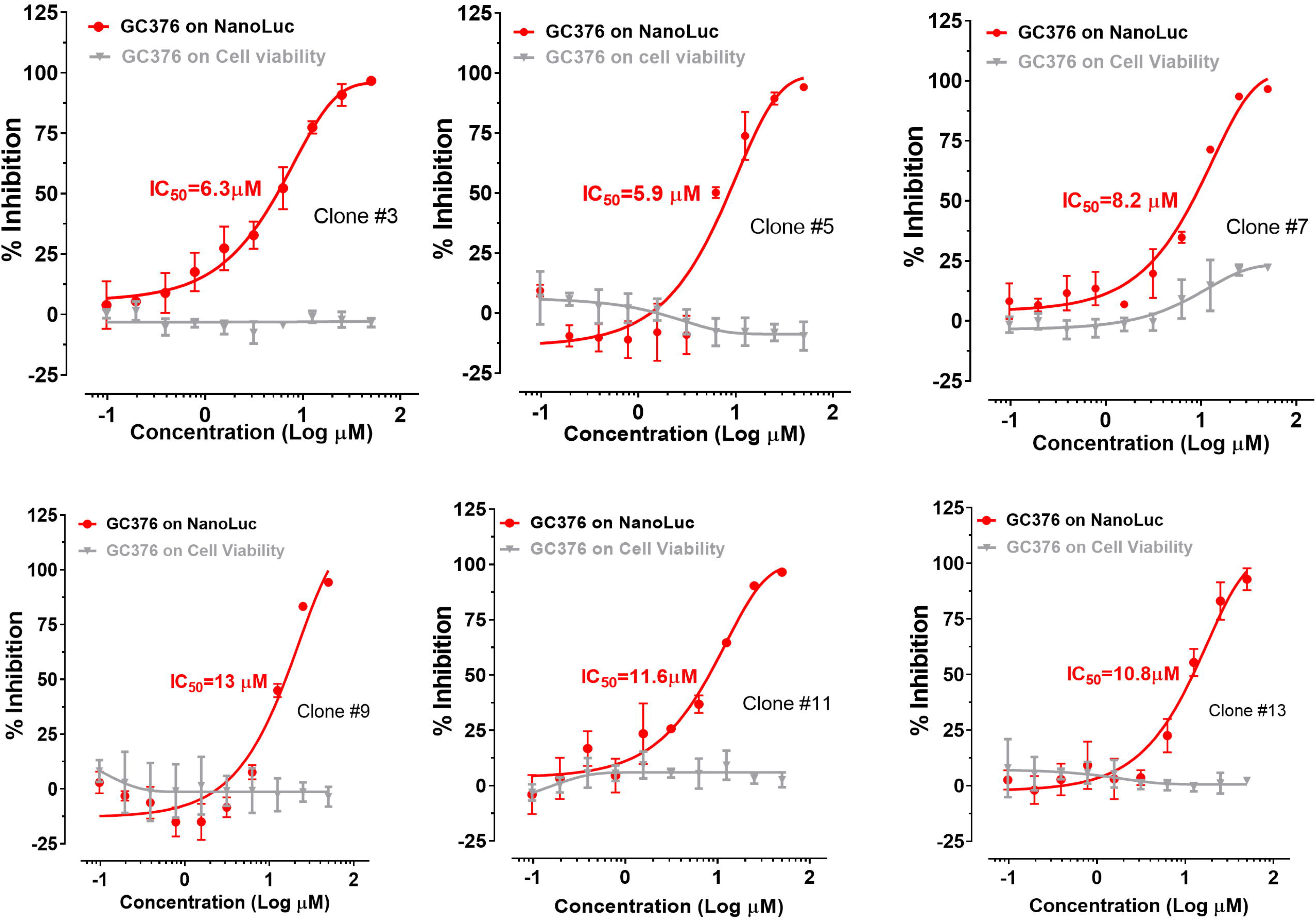
Clonal response to the 3CL protease inhibitor GC376. The half maximal inhibitory concentration (IC50) of GC376 was determined on six stable replicon clones (#3, 5, 7, 9, 11, 13) (red). The effect of GC376 on cell viability (in grey) was simultaneously determined using the Cell Titer-Glo assay.

## DISCUSSION

Two important observations were made in our study regarding establishing stable cell clones harboring SARS-CoV-2 replicon. First, stable cell clones harboring SARS-CoV-2 replicon were obtained when K164A/H165A mutations were introduced to Nsp1. This result is consistent with the structural analysis of Nsp1, which predicts that K164A/H165A reduce the interaction between C-terminus of Nsp1 and the ribosome and hence increase the accessibility of ribosomes to host mRNA. By contrast, R124S/K125E mutations may reduce the binding of the Nsp1 N-terminus to the 5’-UTR of viral mRNA, leaving the C-terminal region of Nsp1 constantly bound to ribosome (15, 20, 21). Consequently, neither viral nor host mRNA can efficiently access the ribosome in the presence of Nsp1^R124S/K125E^. The structural modeling of N128S/K129E could not be performed due to lack of published structure. N128S/K129E, despite being reported to attenuate SARS-CoV Nsp1-mediated inhibition of interferons (22), failed to derive viable cells in a replicon. Secondly, viable cells were only recoverable from the BHK-21 background, demonstrating that SARS-CoV-2-Rep-NanoLuc-Neo-Nsp1^K164A/H165A^ remains cytopathic in other cell types. In addition to Nsp1, other viral factors have been reported to cause cytopathic effect. For example, SARS-CoV-2 PLpro displayed cytopathogenic effect in Vero E6 cells (23). Notably, as this manuscript was in preparation, a similar but non-identical SARS-CoV-2 replicon bearing Nsp1^K164A/H165A^ reportedly failed to generate stable cell clones in Huh-7.5 and BHK-21 cells (24), raising an interesting question whether the detected additional mutations from our stable cell clones facilitated non-cytopathic replication.

In summary, we have established a robust replicon system for future genetic and functional analyses of SARS-CoV-2 replication *in vitro*. To the best of our knowledge, this is the first time that stable cell clones harboring a SARS-CoV-2 replicon could be derived. These clones offer significant advantages over existing replicon systems in that cells may now be conveniently cultured, banked, qualified, and used in a standard BSL-2 laboratory for high throughput screen of compound libraries. This innovation will undoubtedly accelerate the pace of developing treatments for COVID-19.

## MATERIALS AND METHODS

### Cell culture and reagents

The human kidney epithelial cell line Lenti-X 293T was purchased from Takara. The human liver cell line Huh7.5.1 was provided by Dr. Francis Chisari (Scripps Research Institute). The baby hamster kidney fibroblast cell line BHK21 (CCL-10), African green monkey kidney epithelial cells (Vero E6; CRL-1586), Caco-2 (HTB-37), Calu-3 (HTB-55) and A549 (CCL-185) were purchased from the American Type Culture Collection. A549-hACE2 (NR-53821) cells were obtained from BEI Resources. All cell lines were maintained in DMEM supplemented with 5% penicillin and streptomycin, and 10% fetal bovine serum (FBS) at 37 °C with 5% CO2. The SARS-CoV-2 Nucleocapsid antibody (40143-MM05) was purchased from Sino Biological. The SARS-CoV-2 Nsp1 antibody (PA5-116941) was purchased from Themo Fisher Scientific. The β-actin antibody (GTX109639) was purchased from Gentex. Secondary antibodies were purchased from LI-COR Bioscience. GC376 Sodium was purchased from Aobious (AOB36447). Remdesivir was purchased from MedChemExpress (HY-104077).

### Plasmid Construction

Doxycycline-inducible expression of SARS-CoV-2 NP was established in Vero E6, Huh7.5.1, and BHK-21 using TripZ-NP plasmid. NP cDNA was subcloned into pTripZ (AgeI/MluI) using the following primers: TripZ-NPf: 5’-ATATAGACCGGTCCACCATGTCTGATAATGGACCCCA- 3’, TripZ-NPr: 5’- ATATAGACGCGTTTAGGCCTGAGTTGAGTCAG-3’.

### Production of SARS-CoV-2 reporter viruses

SARS-CoV-2 recombinant virus could be generated using a 7-plasmid reverse genetic system which was based on the virus strain (2019-nCoV/USA_WA1/2020) isolated from the first reported SARS-CoV-2 case in the U.S. (10). The initial 7 plasmids were generous gifts from Dr. P-Y Shi (UTMB). Upon receipt, fragment 4 was subsequently subcloned into a low-copy plasmid pSMART LCAmp (Lucigen) to increase stability. Standard molecular biology technique was employed to create the SARS-CoV-2 nanoluciferase and firefly reporter viruses. *In vitro* transcription and electroporation were carried following procedures that were detailed elsewhere (25).

### SARS-CoV-2 Replicon

The SARS-CoV-2-Rep-NanoLuc-Neo replicon was constructed based on the full-length SARS-CoV-2 cDNA infectious clone by replacing the S gene with a nano luciferase gene, and by replacing M and E genes with a neomycin phosphotransferase (Neo) gene. To introduce Nsp1 R124S/K125E, N128S/K129E and K164A/H165A mutations into the SARS-CoV-2-Rep-NanoLuc-Neo replicon, puc57-CoV2-F1 plasmids containing mutated Nsp1 were first created by using overlap PCR method with the following primers:

M13F: GTAAAACGACGGCCAGT
R124S/K125Ef: caaggttcttcttTCGgagaacggtaataaaggagct
R124S/K125Er: ttattaccgttctcCGAaagaagaaccttgcggtaag
N128S/K129Ef: taagaacggtAGTGAGggagctggtggccatagtta
N128S/K129E r: caccagctccCTCACTaccgttcttacgaagaagaa
K164A/H165Af: aaaactggaacactGCcGCcagcagtggtgttacccgtga
K164A/H165Ar: gggtaacaccactgctgGCgGCagtgttccagttttcttgaa
NheIr: cacgagcagcctctgatgca

PCR fragments were digested by Bgl II/Nhe I and ligated into Bgl II/Nhe I digested F1 plasmid. The resulted plasmids were validated by restriction enzyme digestion and Sanger sequencing. Assembly of the 7 plasmids into the replicon and *in vitro* transcription were performed following a published protocol (25).

### RNA Electroporation

Forty-eight hours post doxycycline treatment, BHK-21-NP^Dox-ON^ cells were washed with phosphate buffered saline (PBS), trypsinized, and resuspended in complete growth medium. Cells were pelleted by centrifugation (1,000 x g for 5 min at 4°C), washed twice with ice-cold DMEM, and resuspended in ice-cold Gene Pulser Electroporation Buffer (Bio-Rad) at 1 x 10^7^ cells/ml. Cells (0.4 ml) were then mixed with 10 μg of replicon RNA and 2 μg NP RNA, placed into 4 mm gap electroporation cuvettes, and electroporated at 270 V, 100 Ω, and 950 μF in a Gene Pulser Xcell Total System (Bio-Rad). To establish stable replicon cells, 200 μg/mL of G418 was added to the media between 24 and 48 hours following electroporation, after which culture medium was changed every 2 to 3 days. Three weeks after G418 selection, the resultant foci were counted. All cells were trypsinized and pooled together in a T-75 flask for expansion. Limiting dilution was subsequently performed to derive single cell clones.

### Quantification of viral RNA

Viral RNA was quantified by reverse-transcription quantitative PCR (RT-qPCR) on a StepOnePlus Real-Time PCR System (Applied Biosystems) using Luna Universal Probe One-Step RT-qPCR Kit (New England Biolabs) with an in-house developed protocol. Primers and probes for qPCR were as follows: ORF1ab forward: 5′- CCCTGTGGGTTTTACACTTAA -3′, reverse: 5′- ACGATTGTGCATCAGCTGA-3′, probe: FAM-CCGTCTGCGGTATGTGGAAAGGTTATGG-BHQ1; NanoLuc gene subgenomic mRNA forward: 5′- CCAACCAACTTTCGATCTCTTG-3′, reverse: 5′- GGACTTGGTCCAGGTTGTAG - 3, probe: FAM-ACGAACAATGGTCTTCACACTCGAAGA –BHQ1; Neomycin phosphotransferase gene subgenomic mRNA forward: 5′- CGATCTCTTGTAGATCTGTTCTCTAAA -3′, reverse: 5′- GCCCAGTCATAGCCGAATAG -3′, probe: FAM-ACAAGATGGATTGCACGCAGGTTC-BHQ1. To generate standard plasmids, the cDNAs of SARS-CoV-2 ORF1ab gene, NanoLuc gene sgmRNA and neomycin phosphotransferase gene sgmRNA were cloned into a pCR2.1-TOPO plasmid respectively. The copy number of replicon RNA was calculated by comparing to a standard curve obtained with serial dilutions of the standard plasmid.

### Immunoblotting

Cells were grown in 24-well plates and lysates were prepared with RIPA buffer (50 mM Tris-HCl [pH 7.4]; 1% NP-40; 0.25% sodium deoxycholate; 150 mM NaCl; 1 mM EDTA; protease inhibitor cocktail (Sigma); 1 mM sodium orthovanadate), and insoluble material was precipitated by brief centrifugation. Lysates were loaded onto 4-20% SDSPAGE gels and transferred to a nitrocellulose membrane (LI-COR, Lincoln, NE), blocked with Intercept (TBS) Blocking Buffer ((LI-COR, Lincoln, NE) for 1 h, and incubated with the primary antibody overnight at 4 C. Membranes were blocked with Odyssey Blocking buffer (LI-COR, Lincoln, NE), followed by incubation with primary antibodies at 1:1000 dilutions. Membranes were washed three times with 1X TBS containing 0.05% Tween20 (v/v), incubated with IRDye secondary antibodies (LI-COR, Lincoln, NE) for 1 h, and washed again to remove unbound antibody. Odyssey CLx (LI-COR Biosystems, Lincoln, NE) was used to detect bound antibody complexes.

### Compound Screen

273 compounds (assembled by TargetMol) were diluted in culture media to a final concentration of 5 μM for initial screen. Approximately 1.5 × 10^4^ replicon cells/well were seeded in 96-well plates in the absence of G418. Twenty-four hours later, cell culture media (without G418) was replaced with media containing 5 μM of compounds or the same volume of diluent DMSO. After incubation at 37 °C for specified periods, cells were assayed for NanoLuc activity using Nano-Glo Luciferase Assay System (N1130, Promega) or cell viability using cellTiter-Glo (G7571, Promega).

For validation, A549-hACE2, Caco-2 or Calu-3 cells were seeded in 96-well plates at a density of 10^4^ cells/well. Twenty-four hours later, cells were infected with SARS2-NanoLuc reporter virus in triplicates at an MOI of 0.05 in culture medium containing compounds. After 24 hours at 37 °C, cells were assayed for NanoLuc activity using Nano-Glo Luciferase Assay System or Luciferase Assay System (E1501, Promega).

### Next-generation sequencing

To prepare sequencing libraries, 100 μl of total RNAs were extracted from 5 × 10^5^ replicon cells using RNeasy mini kit (Qiagen, Gaithersburg, MD). 2□μl of each sample was used to assess RNA quality using Agilent 2100 Bioanalyzer (Agilent Technologies, Santa Clara, CA), and the RNA integration numbers (RIN) were all greater than 9. A total of 300 ng of total RNA was used to prepare the sequencing library using Illumina Stranded Total RNA Prep, Ligation with Ribo-Zero Plus. After ribosomal RNA removal, adaptor ligation, and cDNA library concentration and normalization, prepared libraries were loaded onto a NextSeq sequencer (Illumina, San Diego, CA) for deep sequencing of paired-end reads of 2×74 cycles. The numbers of reads mapped to the constructed virus genome range between 27K and 544K for individual samples. Variant calling was performed using Qiagen CLC Genomics Workbench V20 low-frequency variant detection with the requirement of significance of□≥ 5% and minimum frequency□of ≥ 20%. For canonical sgmRNA identification, a set of six sequences were constructed based on the replicon genome, each consisting of 49 nucleotides upstream of the 6-bp transcription regulatory sequence (TRS) motif (ACGAAC) found in the Leader sequence, the TRS-B sequence and 50 nucleotides downstream of each TRS-B, which extend into the coding region of each ORF. The specific sequences covering the junctions are shown as followings:

sgmNanoluc: CAGGTAACAAACCAACCAACTTTCGATCTCTTGTAGATCTGTTCTCTAAACGAACAATGGTCTTCACACTCGAAGATTTCGTTGGGGACTGGCGACAGACAGCCGG (Green, leader sequence; red, TRS-B; orange, NanoLuc).
sgmORF3a: CAGGTAACAAACCAACCAACTTTCGATCTCTTGTAGATCTGTTCTCTAAACGAACTTATGGATTTGTTTATGAGAATCTTCACAATTGGAACTGTAACTTTGAAGCA (Green, leader sequence; red, TRS-B; orange, ORF3a).
sgmNeoR: CAGGTAACAAACCAACCAACTTTCGATCTCTTGTAGATCTGTTCTCTAAACGAACTTATGATTGAACAAGATGGATTGCACGCAGGTTCTCCGGCCGCTTGGGTGGA (Green, leader sequence; red, TRS-B; orange, NeoR)
sgmORF7: CAGGTAACAAACCAACCAACTTTCGATCTCTTGTAGATCTGTTCTCTAAACGACGAACATGAAAATTATTCTTTTCTTGGCACTGATAACACTCGCTACTTGTGAGCT (Green, leader sequence; red, TRS-B; orange, ORF7).
sgmORF8: CAGGTAACAAACCAACCAACTTTCGATCTCTTGTAGATCTGTTCTCTAAACGAACATGAAATTTCTTGTTTTCTTAGGAATCATCACAACTGTAGCTGCATTTCA (Green, leader sequence; red, TRS-B; orange, ORF8).
sgmNP: CAGGTAACAAACCAACCAACTTTCGATCTCTTGTAGATCTGTTCTCTAAACGAACAAACTAAAATGTCTGATAATGGACCCCAAAATCAGCGAAATGCACCCCGCATTACGTT (Green, leader sequence; red, TRS-B; orange, NP).

Short, paired-end reads of RNA samples from 12 stable replicon cell clones were uploaded and analyzed on the NGS platform High performance Integrated Virtual Environment (HIVE)(26). The reads were indexed, deduplicated, and quality metrics were collected upon data ingestion, which verified the high quality of the reads (Fig. S7). Alignment of the reads were performed with HIVE’s native (27) against the seven reference sequences. HIVE Hexagon default parameters, tuned for viral analysis allowing for small indels and mutations, were used and a threshold of 65 bases or longer of the aligned query was applied. Given the fact that the length of the reads was 70bp and the way the subject sequences were constructed, the alignments returned only split reads with patterns from both sides of the investigated junctions.

### MD simulations

We carried out all-atom MD simulations for the complex of nsp1 and the fragment of rRNA (Charcoal gray in Fig. 1B) using the NAMD2.13 package (28) running on the IBM Power Cluster. The atomic coordinates for the complex (a bound state) were obtained from the crystal structure (PDB code: 7K5I) (20). The complex was further solvated in a cubic water box that measures about 78×78×78 Å^3^. Na^+^ and Cl^−^ were added to neutralize the entire simulation system and set the ion concentration to be 0.15 M (Fig. S2A). The final simulation system comprises 47,971 atoms. The built system was first minimized for 10 ps and then equilibrated for 1000 ps in the NPT ensemble (*P* ∼ 1 bar and *T* ∼ 300 K), with atoms in the backbones (of both nsp1 and RNA) harmonically restrained (spring constant *k*=1 kcal/mol/Å^2^). The production run (∼200 ns) was performed in the NPT ensemble, when only constraining the terminals of nsp1 (both N- and C-terminals) and rRNA (both 5’ and 3’-terminals). The same approach was applied in the production run for nsp1 in a 0.15 M NaCl electrolyte (Fig. S2B), a free state required in the free energy perturbation (FEP) calculations (see below). The water box for the nsp1-only simulation also measures about 78×78×78 Å (29). Note that the similar system size for the bound and free states are required for free energy perturbation calculations for mutations with a net charge change.

We used the CHARMM36m force field (29) for proteins and rRNA, the TIP3P model for water (30, 31), the standard force field (32) for Na^+^ and Cl^−^. The periodic boundary conditions (PBC) were applied in all three dimensions. Long-range Coulomb interactions were computed using particle-mesh Ewald (PME) full electrostatics with the grid size of about 1 Å in each dimension. The pair-wise van der Waals (vdW) energies were calculated using a smooth (10-12 Å) cutoff. The temperature *T* was kept at 300 K by applying the Langevin thermostat (33), while the pressure was maintained constant at 1 bar using the Nosé-Hoover method (34). With the SETTLE algorithm (35) enabled to keep all bonds rigid, the simulation time-step was 2 fs for bonded and non-bonded (including vdW, angle, improper and dihedral) interactions, and the time-step for Coulomb interactions was 4 fs, with the multiple time-step algorithm (36).

### Free energy perturbation calculations

After equilibrating the structures in bound and free states, we performed free energy perturbation (FEP) calculations (37). In the perturbation method, many intermediate stages (denoted by λ) whose Hamiltonian *H*(λ)=λ*H*_*f*_ +(1-λ)*H*_*i*_ are inserted between the initial (*H*_*i*_) and final (*H*_*f*_) states to yield a high accuracy. With the softcore potential enabled, λ in each FEP calculation for the bound or free state varies from 0 to 1.0 in 20 perturbation windows (lasting 300 ps in each window), yielding gradual and progressive annihilation and exnihilation processes for mutations at residue 164 (K to A), 165 (H to A), 167 (S to A), 171 (R to A) and 175 (R to A), respectively. In FEP runs for the K164A mutation, the net charge of the MD system changed from 0 to −1 e (where e is the elementary charge). It is important to have similar sizes of the simulation systems for the free and the bound states (38, 39) so that the energy shifts from the Ewald summation (due to the net charge in the final simulation system) approximately cancel out when calculating ∆∆*G*. The same approaches were applied to investigate mutations of R171A and R175A. More detailed procedures can be found in our previous work (40, 41).

### *In Vitro* Cytotoxicity Assay and CC_50_ Determination

Cytotoxicity was determined by cell viability assay as previously described. In brief, the cell viability was measured using Cell-Titer Glo (Promega) according to the manufacturers’ instructions, and luminescence signals were measured by GloMax luminometer. CC_50_ values were calculated using a nonlinear regression curve fit in Prism Software version 9 (GraphPad). The reported CC_50_ values were the results of at least 3 biological or technical replicates.

## Acknowledgments

The authors are grateful to Dr. P-Y Shi (UTMB) who generously shared the 7-plasmid SARS-CoV-2 reverse genetics system. We thank Drs. K. Karagiannis and Santana-Quintero for sgmRNA analysis and Drs. K. Peden, S. Tang, and C.B. Stauft for critical reading of the manuscript. The content of this publication does not reflect the views or policies of the Department of Health and Human Services, nor does mention of trade names, commercial products, or organizations imply endorsement by the U.S. Government.

## References

1. Kim D, Lee JY, Yang JS, Kim JW, Kim VN, Chang H. 2020. The Architecture of SARS-CoV-2 Transcriptome. Cell 181:914–921 e10.

2. Rashid F, Dzakah EE, Wang H, Tang S. 2021. The ORF8 protein of SARS-CoV-2 induced endoplasmic reticulum stress and mediated immune evasion by antagonizing production of interferon beta. Virus Res 296:198350.

3. Xia H, Cao Z, Xie X, Zhang X, Chen JY, Wang H, Menachery VD, Rajsbaum R, Shi PY. 2020. Evasion of Type I Interferon by SARS-CoV-2. Cell Rep 33:108234.

4. Lei X, Dong X, Ma R, Wang W, Xiao X, Tian Z, Wang C, Wang Y, Li L, Ren L, Guo F, Zhao Z, Zhou Z, Xiang Z, Wang J. 2020. Activation and evasion of type I interferon responses by SARS-CoV-2. Nat Commun 11:3810.

5. He X, Quan S, Xu M, Rodriguez S, Goh SL, Wei J, Fridman A, Koeplinger KA, Carroll SS, Grobler JA, Espeseth AS, Olsen DB, Hazuda DJ, Wang D. 2021. Generation of SARS-CoV-2 reporter replicon for high-throughput antiviral screening and testing. Proc Natl Acad Sci U S A 118.

6. Kotaki T, Xie X, Shi PY, Kameoka M. 2021. A PCR amplicon-based SARS-CoV-2 replicon for antiviral evaluation. Sci Rep 11:2229.

7. Wang B, Zhang C, Lei X, Ren L, Zhao Z, Wang J, Huang H. 2021. Construction of Non-infectious SARS-CoV-2 Replicons and Their Application in Drug Evaluation. Virol Sin doi:10.1007/s12250-021-00369-9.

8. Zhang X, Liu Y, Liu J, Bailey AL, Plante KS, Plante JA, Zou J, Xia H, Bopp NE, Aguilar PV, Ren P, Menachery VD, Diamond MS, Weaver SC, Xie X, Shi PY. 2021. A trans-complementation system for SARS-CoV-2 recapitulates authentic viral replication without virulence. Cell 184:2229–2238 e13.

9. Thi Nhu Thao T, Labroussaa F, Ebert N, V’Kovski P, Stalder H, Portmann J, Kelly J, Steiner S, Holwerda M, Kratzel A, Gultom M, Schmied K, Laloli L, Husser L, Wider M, Pfaender S, Hirt D, Cippa V, Crespo-Pomar S, Schroder S, Muth D, Niemeyer D, Corman VM, Muller MA, Drosten C, Dijkman R, Jores J, Thiel V. 2020. Rapid reconstruction of SARS-CoV-2 using a synthetic genomics platform. Nature 582:561–565.

10. Xie X, Muruato A, Lokugamage KG, Narayanan K, Zhang X, Zou J, Liu J, Schindewolf C, Bopp NE, Aguilar PV, Plante KS, Weaver SC, Makino S, LeDuc JW, Menachery VD, Shi PY. 2020. An Infectious cDNA Clone of SARS-CoV-2. Cell Host Microbe 27:841–848 e3.

11. Hou YJ, Okuda K, Edwards CE, Martinez DR, Asakura T, Dinnon KH, 3rd, Kato T, Lee RE, Yount BL, Mascenik TM, Chen G, Olivier KN, Ghio A, Tse LV, Leist SR, Gralinski LE, Schafer A, Dang H, Gilmore R, Nakano S, Sun L, Fulcher ML, Livraghi-Butrico A, Nicely NI, Cameron M, Cameron C, Kelvin DJ, de Silva A, Margolis DM, Markmann A, Bartelt L, Zumwalt R, Martinez FJ, Salvatore SP, Borczuk A, Tata PR, Sontake V, Kimple A, Jaspers I, O’Neal WK, Randell SH, Boucher RC, Baric RS. 2020. SARS-CoV-2 Reverse Genetics Reveals a Variable Infection Gradient in the Respiratory Tract. Cell 182:429–446 e14.

12. Finkel Y, Gluck A, Nachshon A, Winkler R, Fisher T, Rozman B, Mizrahi O, Lubelsky Y, Zuckerman B, Slobodin B, Yahalom-Ronen Y, Tamir H, Ulitsky I, Israely T, Paran N, Schwartz M, Stern-Ginossar N. 2021. SARS-CoV-2 uses a multipronged strategy to impede host protein synthesis. Nature 594:240–245.

13. Kamitani W, Huang C, Narayanan K, Lokugamage KG, Makino S. 2009. A two-pronged strategy to suppress host protein synthesis by SARS coronavirus Nsp1 protein. Nat Struct Mol Biol 16:1134–40.

14. Lokugamage KG, Narayanan K, Nakagawa K, Terasaki K, Ramirez SI, Tseng CT, Makino S. 2015. Middle East Respiratory Syndrome Coronavirus nsp1 Inhibits Host Gene Expression by Selectively Targeting mRNAs Transcribed in the Nucleus while Sparing mRNAs of Cytoplasmic Origin. J Virol 89:10970–81.

15. Yuan S, Peng L, Park JJ, Hu Y, Devarkar SC, Dong MB, Shen Q, Wu S, Chen S, Lomakin IB, Xiong Y. 2020. Nonstructural Protein 1 of SARS-CoV-2 Is a Potent Pathogenicity Factor Redirecting Host Protein Synthesis Machinery toward Viral RNA. Mol Cell 80:1055–1066 e6.

16. Schubert K, Karousis ED, Jomaa A, Scaiola A, Echeverria B, Gurzeler LA, Leibundgut M, Thiel V, Muhlemann O, Ban N. 2020. SARS-CoV-2 Nsp1 binds the ribosomal mRNA channel to inhibit translation. Nat Struct Mol Biol 27:959–966.

17. Lapointe CP, Grosely R, Johnson AG, Wang J, Fernandez IS, Puglisi JD. 2021. Dynamic competition between SARS-CoV-2 NSP1 and mRNA on the human ribosome inhibits translation initiation. Proc Natl Acad Sci U S A 118.

18. Finkel Y, Mizrahi O, Nachshon A, Weingarten-Gabbay S, Morgenstern D, Yahalom-Ronen Y, Tamir H, Achdout H, Stein D, Israeli O, Beth-Din A, Melamed S, Weiss S, Israely T, Paran N, Schwartz M, Stern-Ginossar N. 2021. The coding capacity of SARS-CoV-2. Nature 589:125–130.

19. Dittmar M, Lee JS, Whig K, Segrist E, Li M, Kamalia B, Castellana L, Ayyanathan K, Cardenas-Diaz FL, Morrisey EE, Truitt R, Yang W, Jurado K, Samby K, Ramage H, Schultz DC, Cherry S. 2021. Drug repurposing screens reveal cell-type-specific entry pathways and FDA-approved drugs active against SARS-Cov-2. Cell Rep 35:108959.

20. Ming Shi LW, Pietro Fontana, Setu Vora, Ying Zhang, Tian-Min Fu, Judy Lieberman, Hao Wu. 2020. SARS-CoV-2 Nsp1 suppresses host but not viral translation through a bipartite mechanism. BioRxiv doi:https://doi.org/10.1101/2020.09.18.302901.

21. Vankadari N, Jeyasankar NN, Lopes WJ. 2020. Structure of the SARS-CoV-2 Nsp1/5’-Untranslated Region Complex and Implications for Potential Therapeutic Targets, a Vaccine, and Virulence. J Phys Chem Lett 11:9659–9668.

22. Wathelet MG, Orr M, Frieman MB, Baric RS. 2007. Severe acute respiratory syndrome coronavirus evades antiviral signaling: role of nsp1 and rational design of an attenuated strain. J Virol 81:11620–33.

23. Fu Z, Huang B, Tang J, Liu S, Liu M, Ye Y, Liu Z, Xiong Y, Zhu W, Cao D, Li J, Niu X, Zhou H, Zhao YJ, Zhang G, Huang H. 2021. The complex structure of GRL0617 and SARS-CoV-2 PLpro reveals a hot spot for antiviral drug discovery. Nat Commun 12:488.

24. Ricardo-Lax I, Luna JM, Thao TTN, Le Pen J, Yu Y, Hoffmann HH, Schneider WM, Razooky BS, Fernandez-Martinez J, Schmidt F, Weisblum Y, Trueb BS, Berenguer Veiga I, Schmied K, Ebert N, Michailidis E, Peace A, Sanchez-Rivera FJ, Lowe SW, Rout MP, Hatziioannou T, Bieniasz PD, Poirier JT, MacDonald MR, Thiel V, Rice CM. 2021. Replication and single-cycle delivery of SARS-CoV-2 replicons. Science doi:10.1126/science.abj8430:eabj8430.

25. Xie X, Lokugamage KG, Zhang X, Vu MN, Muruato AE, Menachery VD, Shi PY. 2021. Engineering SARS-CoV-2 using a reverse genetic system. Nat Protoc 16:1761–1784.

26. Simonyan V, Chumakov K, Dingerdissen H, Faison W, Goldweber S, Golikov A, Gulzar N, Karagiannis K, Vinh Nguyen Lam P, Maudru T, Muravitskaja O, Osipova E, Pan Y, Pschenichnov A, Rostovtsev A, Santana-Quintero L, Smith K, Thompson EE, Tkachenko V, Torcivia-Rodriguez J, Voskanian A, Wan Q, Wang J, Wu TJ, Wilson C, Mazumder R. 2016. High-performance integrated virtual environment (HIVE): a robust infrastructure for next-generation sequence data analysis. Database (Oxford) 2016.

27. Santana-Quintero L, Dingerdissen H, Thierry-Mieg J, Mazumder R, Simonyan V. 2014. HIVE-hexagon: high-performance, parallelized sequence alignment for next-generation sequencing data analysis. PLoS One 9:e99033.

28. Phillips JC, Braun R, Wang W, Gumbart J, Tajkhorshid E, Villa E, Chipot C, Skeel RD, Kale L, Schulten K. 2005. Scalable molecular dynamics with NAMD. J Comput Chem 26:1781–802.

29. Huang J, Rauscher S, Nawrocki G, Ran T, Feig M, de Groot BL, Grubmuller H, MacKerell AD, Jr. 2017. CHARMM36m: an improved force field for folded and intrinsically disordered proteins. Nat Methods 14:71–73.

30. William L. Jorgensen JC, Jeffry D. Madura, Roger W. Impey and Michael L. Klein 1983. Comparison of Simple Potential Functions for Simulating Liquid Water. J Chem Phys 79.

31. Neria EF, S.; Karplus, M. 1996. Simulation of Activation Free Energies in Molecular Systems. J Chem Phys 105:1902–1921.

32. Beglov DR, B. 1994. Finite representation of an infinite bulk system: Solvent boundary potential for computer simulations. J Chem Phys 100:9050–9063.

33. Allen MPT, D. J. 1987. Computer Simulation of Liquids. Oxford University Press: New York.

34. Martyna GJT, D. J.; Klein, M. L. 1994. Constant Pressure Molecular Dynamics Algorithms. J Chem Phys 101:4177–4189.

35. Miyamoto SK, P. A. 1992. SETTLE: An Analytical Version of the SHAKE and RATTLE Algorithm for Rigid Water Molecules. J Comp Chem 13:952–962.

36. Tuckerman MB, B. J.; Martyna, G. J. 1992. Reversible multiple time scale molecular dynamics. J Chem Phys 97:1990–2001.

37. Chipot CP, A. 2007. Free energy calculations. Springer.

38. Gerhard Hummer LRP, and Angel E. García. 1996. Free Energy of Ionic Hydration. J Phys Chem 100:1206–1215.

39. Luan B, Chen KL, Zhou R. 2016. Mechanism of Divalent-Ion-Induced Charge Inversion of Bacterial Membranes. J Phys Chem Lett 7:2434–8.

40. Luan B, Huynh T. 2020. In Silico Antibody Mutagenesis for Optimizing Its Binding to Spike Protein of Severe Acute Respiratory Syndrome Coronavirus 2. J Phys Chem Lett 11:9781–9787.

41. Luan B, Huynh T. 2021. Insights into SARS-CoV-2’s Mutations for Evading Human Antibodies: Sacrifice and Survival. J Med Chem doi:10.1021/acs.jmedchem.1c00311.

