## Supplemental Figures and Tables for "Stable Cell Clones Harboring Self-Replicating SARS-CoV-2 RNAs for Drug Screen"

**This PDF file includes:**

Supplemental Figures and Legends

Figs. S1 to S7

Tables S1 -S3

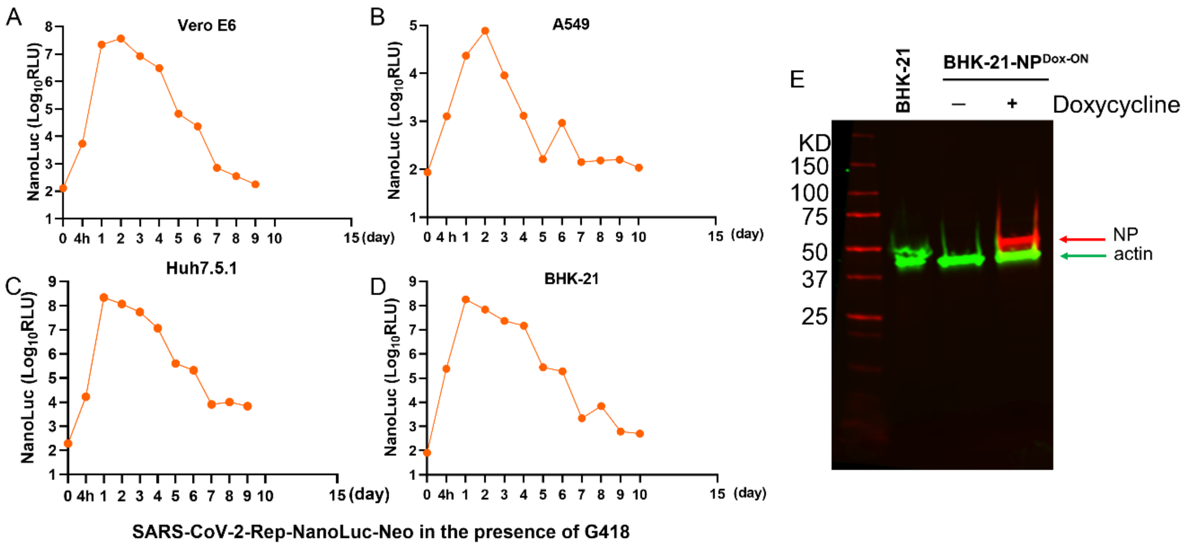


**Fig. S1**. Replication kinetics of SARS-CoV-2-Rep-NanoLuc-Neo in different cell lines. Vero E6 (**A**), A549 (**B**), Huh7.5.1 (**C**) and BHK-21 cells (**D**) were electroporated with replicon RNA. Nano luciferase was measured at indicated time points post-electroporation. (**E**) Generation of BHK21 stable cells that express NP in a doxycycline-inducible manner. Cells were induced with 0.5µg/ml doxycycline and lysed at 48 h postinduction for Western blotting with anti-NP and anti-actin antibodies. Numbers on the left refer to the positions of marker proteins that are given in kilodalton (kDa).

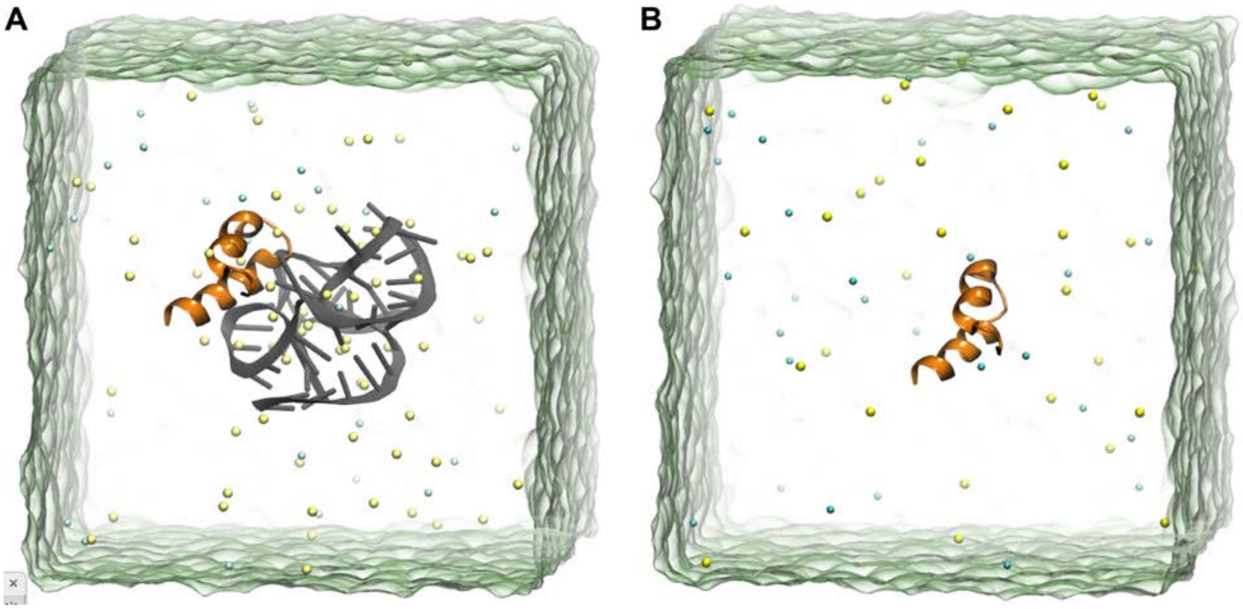


**Fig. S2**. Illustration of MD systems. (**A**) The Nsp-1 and rRNA (fragment) complex in a 0.15 M NaCl electrolyte. The equilibrated structure was used for the FEP calculations of the bound state. (**B**) The Nsp-1 only in a 0.15 M NaCl electrolyte. The equilibrated structure was used for the FEP calculations of the free state.

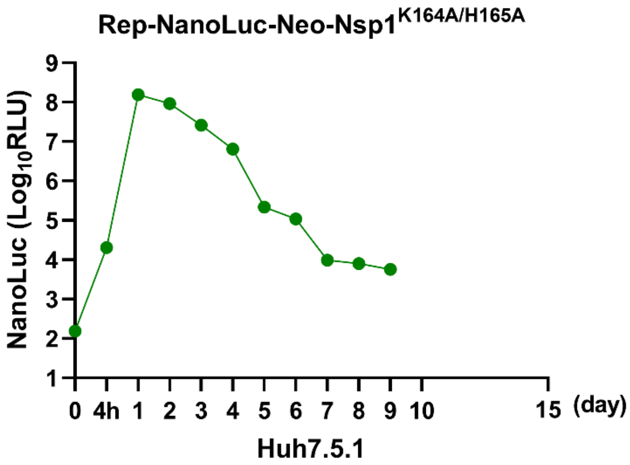


**Fig. S3.** Nanoluciferase kinetics of Rep-NanoLuc-Neo-Nsp1$^{K164A/H165A}$ in Huh7.5.1 cells. Electroporated cells were lysed at indicated time points post-transfection for nanoluciferase quantification.

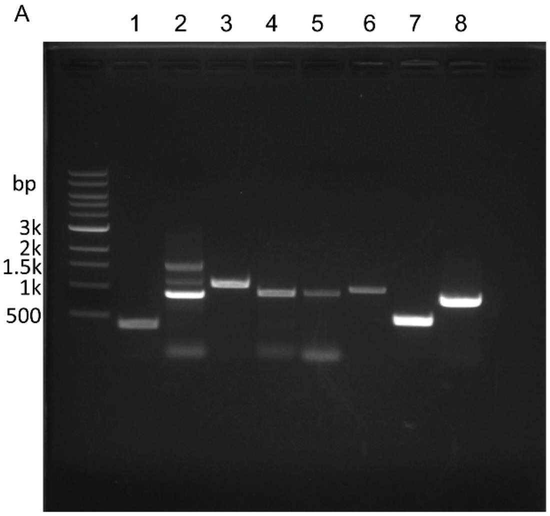


| Lane | Primers | | Amplicon Size (bp) |
|---|---|---|---|
| | Primer name | Sequence | |
| 1 | 32f | ccaaccaactttcgatctcttg | 403 |
| | 434r | caacttctactaagccacaa | |
| 2 | 1000f | agagctatgaattgcagaca | 888 |
| | 1892r | cacgagcagcctctgatgca | |
| 3 | 3000f | ctggtgagtttaaattggct | 1095 |
| | 4072r | tgtcaatgtcactaacaaga | |
| 4 | 7000f | accgctgctttaggtgtttt | 963 |
| | 7965r | ttgacacataagctgactgtagtaa | |
| 5 | 8000f | atgttggtgatagtgcggaa | 960 |
| | 8932r | gtgtgtaacagatgttaccaactgcacta | |
| 6 | 17000f | actcaatatctcagatgagt | 1010 |
| | 17990r | tttcaagacttgtaaattgc | |
| 7 | sgNanof | ccaaccaactttcgatctcttg | 526 |
| | Nanor | tcactccgttgatggttact | |
| 8 | sgEf | cgatctcttgtagatctgttct | 825 |
| | Neor | aagaactcgtcaagaaggcga | |

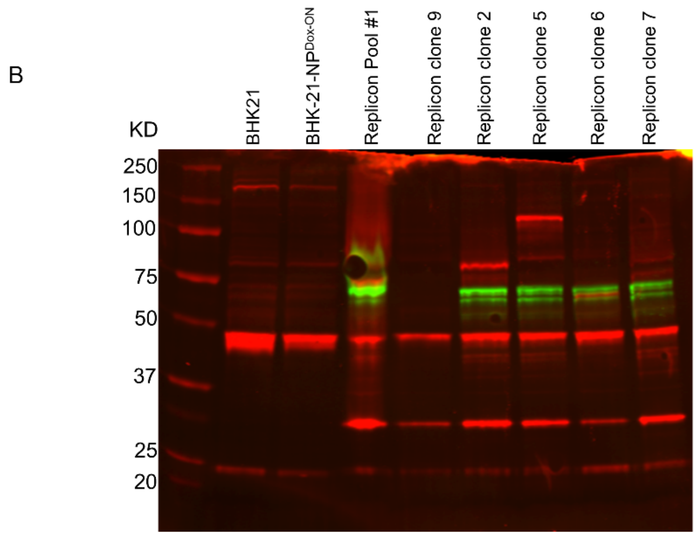


**Fig. S4.** Detection of viral RNA and proteins in replicon cells. (**A**) RT-PCR analysis of viral RNA from replicon cells. The corresponding primer pairs are shown in the right table. The lengths of DNA fragments are indicated in the table. Fragments detected in Lane 1-6 are from different regions of ORF1ab; fragments in lane 7 and 8 represent subgenomic RNAs of the NanoLuc and E (NeoR in relicon). (**B**) Detection of Nsp1 (green) and NP (red) in stable cell clones harboring Rep-NanoLuc-Neo-Nsp1$^{K164A/H165A}$.

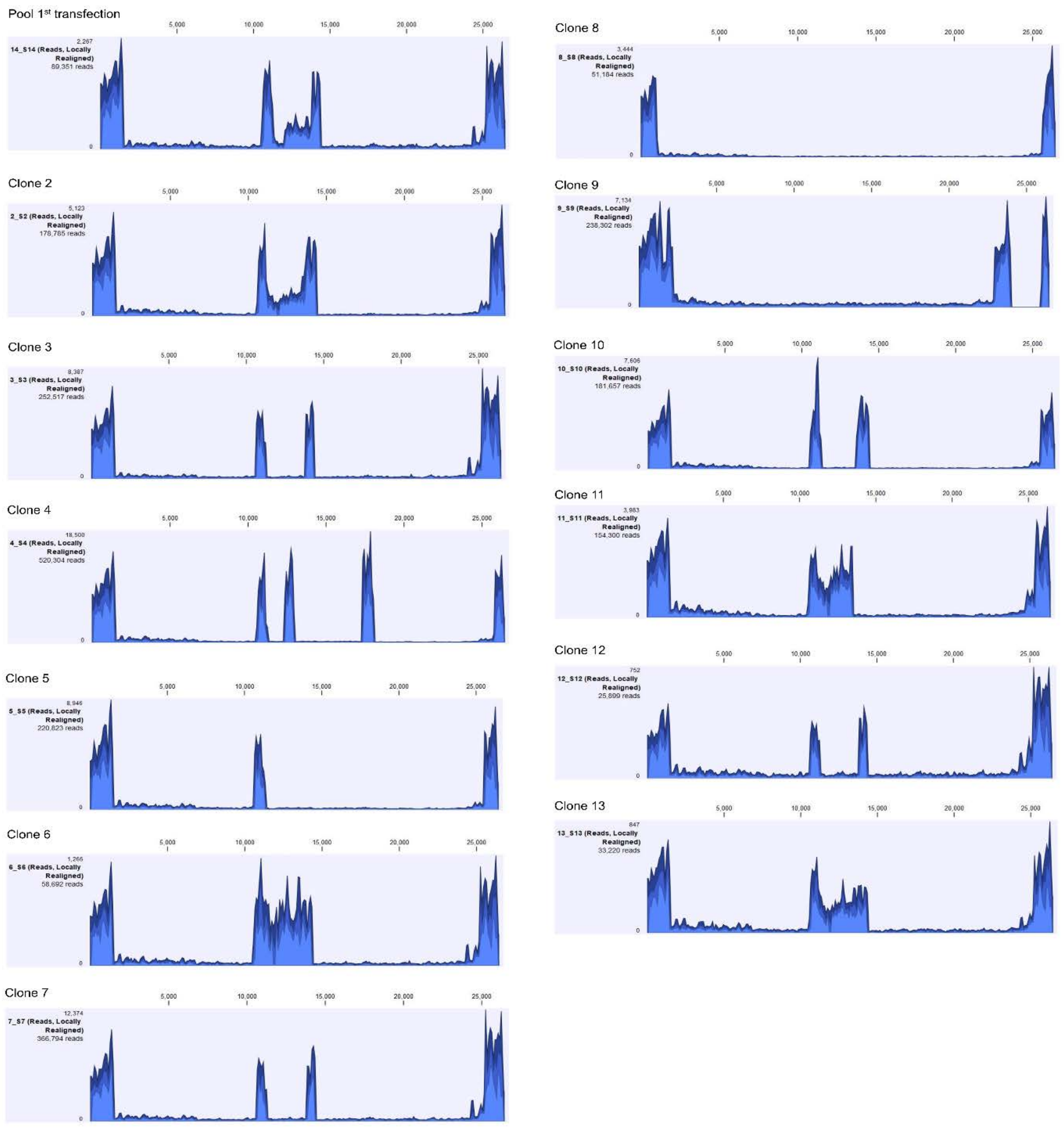


**Fig. S5**. Detection of replicon RNA in stable cell clones. Sequence coverages of the gRNA in each individual stable cell clone as well as in Pool #1 cells were shown. Clone #9 has a truncation which removed the entire ORF7a/b, ORF8 and the first 392 amino acids of the NP region.

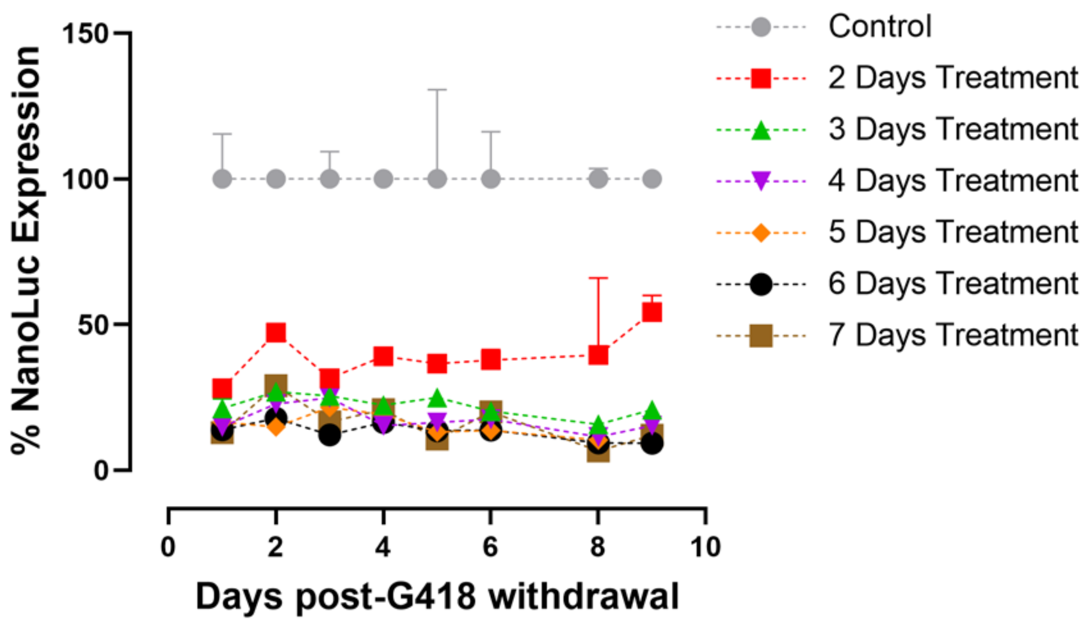


**Fig. S6.** Robust response to Remdesivir in replicon cells. 5 µM Remdesivir was added to Pool #1 cells at different days after G418 withdrawal (day 1 to day 9). Cells were incubated for two to seven days before luciferase was quantified.

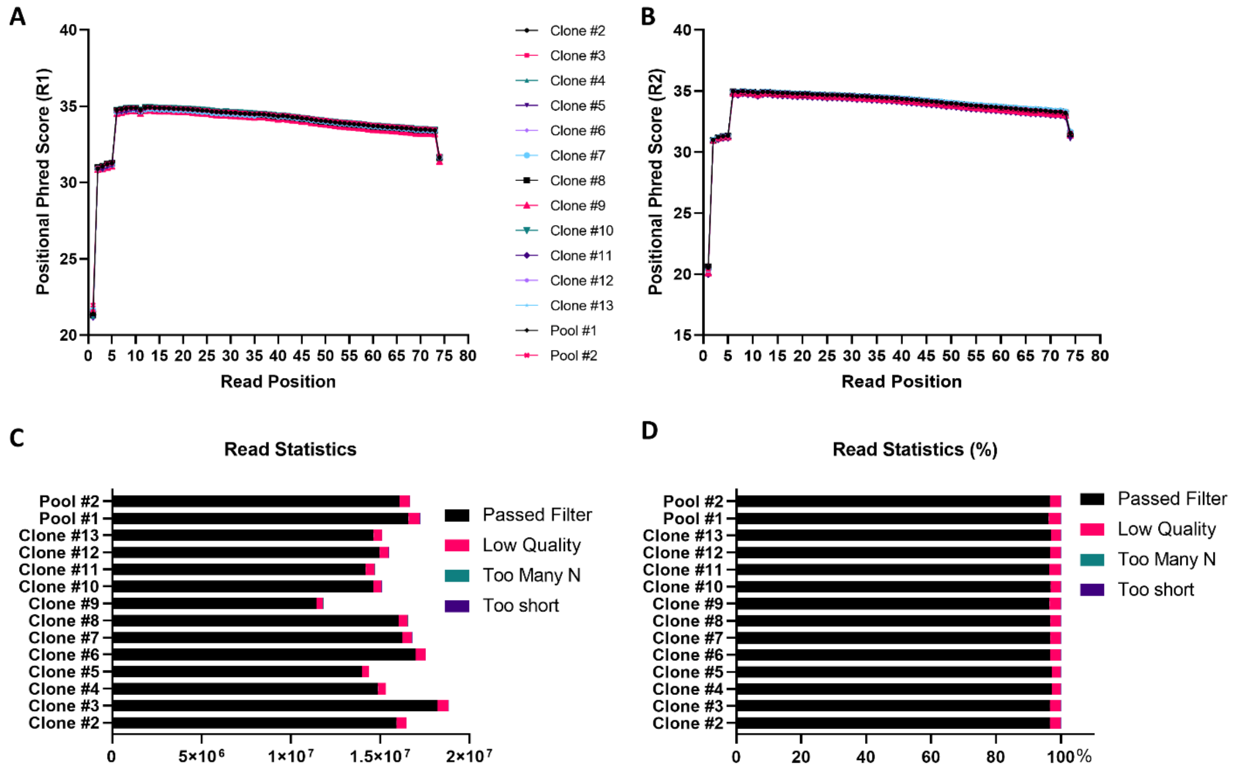


**Fig. S7.** Quality control analysis of sequencing reads. The average Phred quality score remained high (>20) across all position and for all the samples and for both R1 (A) and R2 (B) ends of the paired-end reads. All samples produced more than 10M reads (C) while maintaining a small percentage of low quality reads (D).

**Table S1.** Vial clone numbers

| | $1^{st}$ transfection | $2^{nd}$ transfection | $3^{rd}$ transfection |
|---|---|---|---|
| SARS-CoV-2-Rep-NanoLuc-Neo | 3*(0) | 0 | 0 |
| SARS-CoV-2-Rep-NanoLuc-Neo-$Nsp1^{K164A/H165A}$ | 132 | 7 | 107 |

*Three clones were initially obtained after G418 selection but found to be devoid of the replicon RNA after deep sequencing. $4\times10^6$ BHK-21-$NP^{Dox\text{-}ON}$ cells were electroporated with 10 µg of each RNA for each transfection.

**Table S2** Summary of NGS results of 12 Stable Cell Clones harboring SARS-CoV-2-Rep-NanoLuc-Neo-$Nsp1^{K164A/H165A}$

**Table S3** 273 Anti-COVID-19 compound library

**Table S2**. Summary of NGS results of 12 Stable Cell Clones harboring SARS-CoV-2-Rep-NanoLuc-Neo-Nsp1$^{K164A/H165A}$

| Clone number | Position in MN985325.1 2019-nCoV/USA_WA1/2020 Sequence | Position in Replicon Reference Sequence | Reported MN985325.1 2019-nCoV/USA_WA1/2020 Sequence | Reported Replicon Reference Sequence | Identified Alternative Base | Amino Acid Change | Frequency |
|---|---|---|---|---|---|---|---|
| Clone 2 | 755-760 | 755-760 | AAACAT | AAACAT | GCCGCC | NSP1 K164A H165A | 96.52 |
| | 7486* | 7486 | A | T | T | NSP3 S1589 Synonymous | 100 |
| | 7489* | 7489 | T | A | A | NSP3 T1590 Synonymous | 100 |
| | 6525 | 6525 | C | C | T | NSP3 T1269I | 71.68 |
| | 9755 | 9755 | C | C | A | NSP4 R401S | 100 |
| | 12786 | 12786 | C | C | T | NSP9 T34I | 40.04 |
| | 14724 | 14724 | C | C | T | Synonymous | 100 |
| | 21599-25381 | Deleted | S gene | Nano-Luc | Nano-Luc | S is replaced with NanoLuc | 100 |
| | 26248-27190 | | E and M Gene | NeoR | NeoR | E and M are replaced with neomycin resistant gene (NeoR) | 100 |
| Clone 3 | 755-760 | 755-760 | AAACAT | AAACAT | GCCGCC | NSP1 K164A H165A | 97.25 |
| | 7486* | 7486 | A | T | T | NSP3 S1589 Synonymous | 100 |
| | 7489* | 7489 | T | A | A | NSP3 T1590 Synonymous | 100 |
| | 9755 | 9755 | C | C | A | NSP4 R401S | 100 |
| | 13356 | 13356 | C | C | T | NSP10 T111I | 100 |
| | 21599-25381 | Deleted | S gene | Nano-Luc | Nano-Luc | S is replaced with NanoLuc | 100 |
| | 26248-27190 | | E and M Gene | NeoR | NeoR | E and M are replaced with NeoR | 100 |
| Clone 4 | 755-760 | 755-760 | AAACAT | AAACAT | GCCGCC | NSP1 K164A H165A | 97.25 |
| | 9755 | 9755 | C | C | A | NSP4 R401S | 100 |
| | 7486* | 7486 | A | T | T | NSP3 S1589 Synonymous | 100 |
| | 7489* | 7489 | T | A | A | NSP3 T1590 Synonymous | 99.42 |
| | 19722 | 19722 | A | A | C | NSP15 K34N | 75.36 |
| | N/A | 21788-21789 | N/A | Insertion | A | Nanoluc E76 | 66.06 |
| | N/A | 21833 | N/A | A | G | Nanoluc K91E | 99.09 |
| | N/A | 21862-21863 | N/A | AA | GG | Nanoluc V100 synonymous I101V | 100 |
| | 21599-25381 | Deleted | S gene | Nano-Luc | Nano-Luc | S is replaced with NanoLuc | 100 |
| | 26248-27190 | | E and M Gene | NeoR | NeoR | E and M are replaced with NeoR | 100 |
| | 28955 | 25497 | A | A | G | NP N228D | 97.46 |
| | 29446 | 25988 | T | T | GA | NP T391 | 90.27 |
| | 29449 | 25991 | G | G | A | NP V392 | 92.21 |
| Clone 5 | 755-760 | 755-760 | AAACAT | AAACAT | GCCGCC | NSP1 K164A H165A | 96.39 |
| | 7486* | 7486 | A | T | T | NSP3 S1589 Synonymous | 100 |
| | 7489* | 7489 | T | A | A | NSP3 T1590 Synonymous | 100 |

| | | | | | | | |
|---|---|---|---|---|---|---|---|
| | 9755 | 9755 | C | C | A | NSP4 R401S | 100 |
| | 15006 | 15006 | G | G | T | NSP12 D523Y | 100 |
| | N/A | 21833 | N/A | A | G | Nanoluc K91E | 100 |
| | N/A | 21862-21863 | N/A | AA | GG | Nanoluc<br>V100 synonymous<br>I101V | 100 |
| | 21599-25381 | Deleted | S gene | Nano-Luc | Nano-Luc | S is replaced with NanoLuc | 100 |
| | 26248-27190 | | E and M Gene | NeoR | NeoR | E and M are replaced with NeoR | 100 |
| | 26078 | 22772 | C | C | T | ORF3a T229I | 100 |
| | 28955 | 25497 | A | A | G | NP N228D | 98.41 |
| | 29448 | 25990 | T | T | A | NP V392E | 78.44 |
| Clone 6 | 755-760 | 755-760 | AAACAT | AAACAT | GCCGCC | NSP1 K164A H165A | 97.46 |
| | 7486* | 7486 | A | T | T | NSP3 S1589 Synonymous | 100 |
| | 7489* | 7489 | T | A | A | NSP3 T1590 Synonymous | 100 |
| | 11075 | 11075 | Deletion T | Deletion T | | NSP6 | 38.16 |
| | 15221 | 15221 | T | T | G | NSP12 F594C | 100 |
| | 21599-25381 | Deleted | S gene | Nano-Luc | Nano-Luc | S is replaced with NanoLuc | 100 |
| | 26248-27190 | | E and M Gene | NeoR | NeoR | E and M are replaced with NeoR | 100 |
| Clone 7 | 755-760 | 755-760 | AAACAT | AAACAT | GCCGCC | NSP1 K164A H165A | 96.83 |
| | 2189 | 2189 | C | C | T | NSP2 L462F | 95.45 |
| | 7486* | 7486 | A | T | T | NSP3 S1589 Synonymous | 99.45 |
| | 7489* | 7489 | T | A | A | NSP3 T1590 Synonymous | 100 |
| | 9755 | 9755 | C | C | A | NSP4 R401S | 99.17 |
| | 12357 | 12357 | C | C | T | NSP8 T89I | 100 |
| | 13356 | 13356 | C | C | T | NSP10 T111I | 100 |
| | 13953 | 13953 | A | A | C | NSP 12 I171 Synonymous | 94.36 |
| | 21599-25381 | Deleted | S gene | Nano-Luc | Nano-Luc | S is replaced with NanoLuc | 100 |
| | 26248-27190 | | E and M Gene | NeoR | NeoR | E and M are replaced with NeoR | 100 |
| | N/A | 23478 | N/A | C | T | NeoR synonymous | 100 |
| Clone 8 | 755-760 | 755-760 | AAACAT | AAACAT | GCCGCC | NSP1 K164A H165A | 97.57 |
| | 1392 | 1392 | C | C | T | NSP2 S196L | 100 |
| | 7486* | 7486 | A | T | T | NSP3 S1589 Synonymous | 100 |
| | 7489* | 7489 | T | A | A | NSP3 T1590 Synonymous | 95.65 |
| | 9755 | 9755 | C | C | A | NSP4 R401S | 90 |
| | 13356 | 13356 | C | C | T | NSP10 T111I | 94.44 |
| | 15521 | 15521 | T | T | G | NSP12 F594C | 100 |
| | 21599-25381 | Deleted | S gene | Nano-Luc | Nano-Luc | S is replaced with NanoLuc | 100 |
| | 26248-27190 | | E and M Gene | NeoR | NeoR | E and M are replaced with NeoR | 100 |
| Clone 9 | 755-760 | 755-760 | AAACAT | AAACAT | GCCGCC | NSP1 K164A H165A | 97.64 |
| | 2416 | 2416 | C | C | T | NSP2 Y537 synonymous | 99.79 |

| | | | | | | | |
|---|---|---|---|---|---|---|---|
| | 6896 | 6896 | C | C | T | NSP3 L1393 synonymous | 98.99 |
| | 7486* | 7486 | A | T | T | NSP3 S1589 Synonymous | 100 |
| | 7489* | 7489 | T | A | A | NSP3 T1590 Synonymous | 100 |
| | 9755 | 9755 | C | C | A | NSP4 R401S | 99.04 |
| | 11286 | 11286 | T | T | G | NSP6 L105W | 98.69 |
| | 13356 | 13356 | C | C | T | NSP10 T111I | 99.28 |
| | 21599-25381 | Deleted | S gene | Nano-Luc | Nano-Luc | S is replaced with NanoLuc | 100 |
| | 26248-27190 | | E and M Gene | NeoR | NeoR | E and M are replaced with NeoR | 100 |
| | 26132 | 22826 | A | A | C | ORF3A H246P | 100 |
| | 27392-29448 | 23934-25990 | ORF7-N | deletion | | Deletion ORF7-N | 100 |
| Clone 10 | 755-760 | 755-760 | AAACAT | AAACAT | GCCGCC | NSP1 K164A H165A | 97.29 |
| | 2751 | 2751 | C | C | T | NSP3 T11I | 100 |
| | 7486* | 7486 | A | T | T | NSP3 S1589 Synonymous | 98.53 |
| | 7489* | 7489 | T | A | A | NSP3 T1590 Synonymous | 98.59 |
| | 8558 | 8558 | A | A | G | NSP4 I2V | 100 |
| | 9755 | 9755 | C | C | A | NSP4 R401S | 100 |
| | 13356 | 13356 | C | C | T | NSP10 T111I | 100 |
| | 21599-25381 | Deleted | S gene | Nano-Luc | Nano-Luc | S is replaced with NanoLuc | 100 |
| | 26248-27190 | | E and M Gene | NeoR | NeoR | E and M are replaced with NeoR | 100 |
| | N/A | 23190 | | C | T | NeoR V84 synonymous | 100 |
| | 27625 | 24167 | C | C | T | ORF7 R78C | 81.40 |
| Clone 11 | 755-760 | 755-760 | AAACAT | AAACAT | GCCGCC | NSP1 K164A H165A | 97.87 |
| | 7486* | 7486 | A | T | T | NSP3 S1589 Synonymous | 100 |
| | 7489* | 7489 | T | A | A | NSP3 T1590 Synonymous | 100 |
| | 9755 | 9755 | C | C | A | NSP4 R401S | 100 |
| | 19102 | 19102 | C | C | T | NSP14 | 100 |
| | 21599-25381 | Deleted | S gene | Nano-Luc | Nano-Luc | S is replaced with NanoLuc | 100 |
| | 26248-27190 | | E and M Gene | NeoR | NeoR | E and M are replaced with NeoR | 100 |
| Clone 12 | 755-760 | 755-760 | AAACAT | AAACAT | GCCGCC | NSP1 K164A H165A | 98.96 |
| | 1182 | 1182 | T | T | C | NSP2 V126A | 20.42 |
| | 2954 | 2954 | T | T | G | NSP3 L79V | 66.67 |
| | 7486* | 7486 | A | T | T | NSP3 S1589 Synonymous | 100 |
| | 7489* | 7489 | T | A | A | NSP3 T1590 Synonymous | 100 |
| | 9755 | 9755 | C | C | A | NSP4 R401S | 100 |
| | 11750 | 11750 | C | C | A | NSP6 L260I | 69.23 |
| | 12357 | 12357 | C | C | T | NSP8 T89I | 100 |
| | 13356 | 13356 | C | C | T | NSP10 T111I | 100 |

| | | | | | | | |
|---|---|---|---|---|---|---|---|
| | 13499 | 13499 | C | C | T | NSP12 T20I | 100 |
| | 13953 | 13953 | A | A | C | NSP 12 I171 Synonymous | 88.15 |
| | 21599-25381 | Deleted | S gene | Nano-Luc | Nano-Luc | S is replaced with NanoLuc | 100 |
| | 26248-27190 | | E and M Gene | NeoR | NeoR | E and M are replaced with NeoR | 100 |
| Clone 13 | 755-760 | 755-760 | AAACAT | AAACAT | GCCGCC | NSP1 K164A H165A | 96.82 |
| | 1392 | 1392 | C | C | T | NSP2 S196L | 24.01 |
| | 7486* | 7486 | A | T | T | NSP3 S1589 Synonymous | 100 |
| | 7489* | 7489 | T | A | A | NSP3 T1590 Synonymous | 100 |
| | 11075 | 11075 | T deletion | T deletion | | NSP6 F35 | 23.86 |
| | 21599-25381 | Deleted | S gene | Nano-Luc | Nano-Luc | S is replaced with NanoLuc | 100 |
| | 26248-27190 | | E and M Gene | NeoR | NeoR | E and M are replaced with NeoR | 100 |

*These two mutations were introduced to differentiate the infectious clone-derived virus from the parental clinical isolate 2019-nCoV/USA_WA1/2020.

**Table S3**. 273 compound library

| ID | Name | Synonyms | CAS | SMILES | Formula | MolWt | Target | Bioactivity | Total_Score |
|---|---|---|---|---|---|---|---|---|---|
| T2934 | Bilirubin | idin;Hemetoidin; | 635-65-4 | (c1CCC(=O | O6 | 584.68 | Endogenous Metabolite | and one of the major end products of | 3CLpro(8.96);nsp16(9.16) |
| T3281 | Delapril Hydrochloride | Hydrochloride;Ali | 83435-67-0 | [C@H](CC | N2O5 | 489.01 | RAAS inhibitor | into two active metabolites, 5-hydroxy | 3CLpro(8.3) |
| T5014 | Prostaglandin E2 (PGE2) | staglandin | 363-24-6 | @H](O)\C= | C20H32O5 | 352.47 | Metabolite | substance that participate in a wide | 3CLpro(8.19) |
| T6109 | Darapladib | SB-480848 | 356057-34-6 | CN(Cc1ccc | N4O2S | 666.77 | Phospholipase inhibitor | substituted pyrimidone with inhibitory | 3CLpro(7.97) |
| T0158 | Mitoxantrone hydrochloride | dihydrochloride; | 70476-82-3 | CNc1c2C(= | N4O6 | 517.4 | Topoisomerase inhibitor | hydrochloride salt of an | 3CLpro(7.71);PLpro(9.22) |
| G211-0291 | G211-0291 | | 959508-41-9 | NCC(=CC1 | O2 | 399.5 | HEK293 | HEK293 inhibitor | 3CLpro(7.57) |
| T0467 | Sildenafil citrate | UK-92480 citrate | 171599-83-0 | (O)(CC(O) | O11S | 666.7 | PDE inhibitor | monophosphate (cGMP)-specific | 3CLpro(7.44) |
| C924-0274 | C924-0274 | | 890825-65-7 | (=O)N/C1= | 3O2 | 421.52 | Cellular tumor antigen p53 | Cellular tumor antigen p53 inhibitor | 3CLpro(7.42) |
| C519-1772 | C519-1772 | | 902574-29-2 | C=2C=C1C | O4S | 544.76 | DNA polymerase beta | DNA polymerase beta inhibitor | 3CLpro(7.32) |
| C697-0280 | C697-0280 | | 902563-73-9 | )C=CN(C)C | O4S | 450.6 | Nonstructural protein 1 | Nonstructural protein 1 inhibitor | 3CLpro(7.32) |
| T2873 | Ginsenoside Rg2 | Rg2;Prosapogeni | 52286-74-5 | 1O[C@@H | 3 | 785.01 | GSK-3 antagonist | active components of ginseng, act as | 3CLpro(7.31) |
| T2310 | CHIR99021 | 99021;CHIR- | 252917-06-9 | ]1)- | N8 | 465.34 | GSK-3 | inhibitor (IC50: 10/6.7 nM). | 3CLpro(7.3) |
| T5S1103 | Isoliensinine | Isoliensinin | 6817-41-0 | [C@H]2N( | O6 | 610.75 | Antioxidant | bisbenzyltetrahydroisoquinoline | 3CLpro(7.27) |
| C738-0291 | C738-0291 | | 902954-26-1 | N(CC(=CC | O4 | 399.49 | G(s), subunit alpha | G(s), subunit alpha inhibitor | 3CLpro(7.21) |
| D072-0267 | D072-0267 | | 855714-75-9 | C=1N=C(O | 2O4S | 402.45 | Ataxin-2 | Ataxin-2 inhibitor | 3CLpro(7.12) |
| 2995-0491 | 2995-0491 | | 312598-19-9 | )(C(C1(C)C | O5S | 431.51 | Beta-lactamase AmpC | Beta-lactamase AmpC inhibitor | 3CLpro(7.04) |
| T6332 | Pevonedistat | MLN4924 | 905579-51-3 | )OC[C@@ | O4S | 443.52 | E1 Activating inhibitor | small molecule NEDD8-activating | 3CLpro(7.02);nsp15(7.47) |
| C115-0510 | C115-0510 | | 685868-67-1 | C@]([H])(C | O7 | 496.57 | Arachidonate 15-lipoxygenase | Arachidonate 15-lipoxygenase inhibitor | 3CLpro(7.01) |
| T4961 | 9'-Methyl lithospermate B | | 1167424-31-8 | C@@H](C | 6 | 732.64 | Others | NA | Domain(11.93) |
| T0772 | Troxerutin | tin | 7085-55-4 | 1O[C@@H | 9 | 742.67 | NOD-like Receptor (NLR) | isolated from Sophora japonica. It has | nsp16(9.84) |
| T7740 | 4 diTFA(245443-52-1(free | | T7740 | )(F)F.OC(= | N8O11 | 894.81 | Protease-activated Receptor | diTFA, 2454 is the proteinase- | nsp16(9.61) |
| T3054 | Daurisoline | (R,R)-Daurisoline | 70553-76-3 | c(c(cc2[C@ | O6 | 610.75 | Autophagy | also an autophagy blocker. | nsp16(9.6) |
| T2S0257 | succinate | | 786593-06-4 | COC(=O)C | 0 | 532.58 | Others | traditional Chinese medicine used in | nsp16(9.53);RdRP(10.83) |
| T5726 | Specneuzhenide | Nuezhenide | 449733-84-0 | 1=CO[C@ | 7 | 686.65 | Others | isolated from Ligustrum sinense. | nsp16(9.04) |
| T1795 | Carfilzomib | PR-171 | 868540-17-4 | C(=O)[C@ | O7 | 719.91 | Proteasome inhibitor | proteasome inhibitor and antineoplastic | nsp16(9.03) |
| T4049 | Genz-123346 free base | | 491833-30-8 | CCCCC)N[ | O4 | 418.57 | Glucokinase inhibitor | synthase that blocks the conversion of | nsp16(8.96) |
| T6130 | Skepinone-L | CBS3830 | 1221485-83-1 | H](O)COc1 | NO4 | 425.42 | p38 MAPK inhibitor | activated protein kinase inhibitor. | nsp16(8.74) |
| T6917 | Oleuropein | | 32619-42-4 | 1=CO[C@ | 3 | 540.51 | Aromatase inhibitor; ROS inhibitor | polyphenol isolated from olive leaf. | nsp16(8.58);RdRP(10.98) |
| T3099 | Pinometostat | EPZ-5676 | 1380288-87-8 | C@H]1O[C | O3 | 562.71 | Histone Methyltransferase inhibitor | studying the treatment of Leukemia, | nsp16(8.56);nsp15(6.96) |
| T0447 | Carvedilol | 105517 | 72956-09-3 | 1OCCNCC( | O4 | 406.47 | modulator; Integrin inhibitor; NADPH | salt form of carvedilol, a racemic | nsp16(8.55);nsp15(7.47) |
| T5841 | Travoprost | isopropyl | 157283-68-6 | O)CCC\C= | O6 | 500.55 | Prostaglandin Receptor | and ocular hypertension,is a potent and | nsp16(8.49) |
| T5234 | Glycoursodeoxycholic acid | glycine;GUDCA | 64480-66-6 | CC(=O)NC | 5 | 449.62 | Others | glycine and a bile acid-glycine | nsp16(8.47) |
| T4678 | Fmoc-Val-Cit-PAB | | 159858-22-7 | H](NC(=O) | O6 | 601.69 | Others | antibody-drug-conjugation (ADC). | nsp16(8.46);RdRP(10.48) |
| C522-0732 | C522-0732 | | 901716-39-0 | CN(CC2)C | N4O3 | 458.99 | Beta-lactamase AmpC | Beta-lactamase AmpC inhibitor | nsp16(8.29) |
| E859-1859 | E859-1859 | | 894561-23-0 | C(C=C2)= | O2 | 484.61 | TAR DNA-binding protein 43 | TAR DNA-binding protein 43 inhibitor | nsp16(8.27);X Domain(10.05) |
| T3579 | PLX8394 | | 1393466-87-9 | CCN(C1)S( | N6O3S | 542.54 | Raf inhibitor | serine/threonine-protein kinase B-Raf | nsp16(8.23);nsp15(7.15) |
| D011-0999 | D011-0999 | | 883644-08-4 | C(C1=CC1) | O4 | 485.59 | factor 2 | factor 2 inhibitor | nsp16(8.2) |
| D126-0074 | D126-0074 | | 901663-71-6 | C(=C2)O)= | O5 | 479.58 | HEK293 | HEK293 inhibitor | nsp16(8.18) |
| C147-0154 | C147-0154 | | 825601-75-0 | C/C=C1/N | O | 371.49 | Cellular tumor antigen p53 | Cellular tumor antigen p53 inhibitor | nsp16(8.16) |
| T2544 | Bazedoxifene acetate | 424;TSE | 198481-33-3 | c1c(cc2c(c | O5 | 530.67 | inhibitor | selective estrogen receptor modulator | nsp16(8.12) |
| 8015-6465 | 8015-6465 | | 696634-21-6 | CC(=O)C1 | O5 | 398.46 | Ataxin-2 | Ataxin-2 inhibitor | nsp16(8.08) |
| T6162 | BS-181 HCl | hydrochloride | 1397219-81-6 | cnn2c(NCc | HCl | 416.99 | CDK inhibitor | inhibitor with IC50 of 21 nM. It is more | nsp16(8.03) |
| T6883 | LY3023414 | GTPL8918 | 1386874-06-1 | c3cc(c4cc( | O3 | 406.48 | PI3K inhibitor | inhibitor of the class I PI3K isoforms, | nsp16(8.02) |
| T6S1768 | Narcissoside | mnetin 3- | 604-80-8 | c1O)- | 6 | 624.54 | Antioxidant | B.flavum flavonoid and rutin, could be | nsp16(8.02) |
| T3217 | PF-CBP1 hydrochloride | PF-CBP1 HCl | 2070014-93-4 | c(cc1)CCc1 | N4O3 | 525.08 | Epigenetic Reader Domain inhibitor | of the bromodomain of CREB-binding | nsp16(8) |
| E589-2554 | E589-2554 | | 892739-01-4 | (C1=O)CC( | O6S2 | 491.59 | Geminin | Geminin inhibitor | PLpro(10.32) |
| G071-0431 | G071-0431 | | 895101-33-4 | C2)CC(C(C | O3S2 | 530.67 | proteolytic subunit | proteolytic subunit inhibitor | PLpro(10.01) |
| 7472-0051 | 7472-0051 | | 892246-53-6 | CC(O1)C)( | 4O2S | 426.52 | Ataxin-2 | Ataxin-2 inhibitor | PLpro(9.89) |
| D074-0222 | D074-0222 | | 879938-29-1 | N=C(C=1[C | O4 | 497.6 | Microtubule-associated protein tau | inhibitor | PLpro(9.83) |
| T5318 | CREB inhibitor | CREB inhibitor | 1433286-70-4 | c1cc2ccccc | N3O5 | 620.52 | Epigenetic Reader Domain | inhibitor (IC50: 81 nM). | PLpro(9.83) |
| T3108 | CUDC101 | 101;CUDC-101 | 1012054-59-9 | cc(c(c2)OC | O4 | 434.49 | inhibitor | HDAC, EGFR and HER2 with IC50s of | PLpro(9.67) |
| T2727 | Salvianolic acid B | | 115939-25-8 | @@H](Cc1 | 6 | 718.59 | Others | pharmaceutical compound present in | PLpro(9.56) |
| G114-0456 | G114-0456 | | 895099-65-7 | (C1=O)CC( | O4S2 | 433.55 | homolog 3 | homolog 3 inhibitor | PLpro(9.45) |
| K788-6101 | K788-6101 | | 899212-06-7 | (=O)NCCC | N5O2 | 488.01 | Beta-lactamase AmpC | Beta-lactamase AmpC inhibitor | PLpro(9.4) |
| G240-0046 | G240-0046 | | 895252-40-1 | 1OC)C=C( | O7S | 448.5 | factor 2 | factor 2 inhibitor | PLpro(9.18) |
| C200-4180 | C200-4180 | | 892305-29-2 | CC(NC(C= | N5O4S | 514.01 | factor 2 | factor 2 inhibitor | PLpro(9.12) |
| K784-0203 | K784-0203 | | 422555-16-6 | (NC(=C(C1 | 2O3S | 430.55 | Prelamin-A/C | Prelamin-A/C inhibitor | PLpro(9.06);nsp15(6.89) |
| G406-0489 | G406-0489 | | 959561-08-1 | CC1)(C1=C | O6S | 487.58 | HEK293 | HEK293 inhibitor | PLpro(8.98) |
| C336-0153 | C336-0153 | | 959510-20-4 | =N[C@@]1 | O4S | 533.65 | Inositol monophosphatase 1 | Inositol monophosphatase 1 inhibitor | PLpro(8.96);X Domain(10.41) |
| K786-5309 | K786-5309 | | 697278-01-6 | @]([H])(CN | O2 | 332.49 | Survival motor neuron protein | Survival motor neuron protein inhibitor | PLpro(8.92) |
| T3952 | TPEN | TPEDA | 16858-02-9 | cccn1)Cc1c | C26H28N6 | 424.55 | Autophagy | heavy metal chelator. | PLpro(8.91) |
| Y050-2147 | Y050-2147 | | 887600-07-9 | N1CCOCC | O6S | 398.48 | factor 2 | factor 2 inhibitor | PLpro(8.91) |
| T3112 | Verteporfin | MA | 129497-78-5 | Cc1c(C)c2c | O8 | 718.79 | VDA inhibitor | derivative monoacid ring A, can inhibit | PLpro(8.9) |
| E946-0756 | E946-0756 | | 950395-17-2 | )C(=NC=2 | O3 | 437.55 | Microtubule-associated protein tau | inhibitor | PLpro(8.88) |
| K784-7502 | K784-7502 | | 688796-67-0 | /C=C(/[Cl]) | N5OS | 437.95 | Cellular tumor antigen p53 | Cellular tumor antigen p53 inhibitor | PLpro(8.88) |
| E642-1065 | E642-1065 | | 893161-16-5 | CC(C=C2F | 4OS | 496.61 | homolog 3 | homolog 3 inhibitor | PLpro(8.83) |
| T4259 | GLPG0187 | | 1320346-97-1 | c1)S(=O)(= | O5S | 595.72 | Integrin | receptor antagonist, inhibits αvβ1- | PLpro(8.81) |
| T3631 | PF8380 | 8380 | 1144035-53-9 | 1CCC(=O)c | N3O5 | 478.33 | PDE inhibitor | available autotaxin inhibitor (IC50: 2.8 | PLpro(8.79) |
| G211-0145 | G211-0145 | | 959490-59-6 | C1=CC=C | O3 | 429.52 | Glucagon-like peptide 1 receptor | inhibitor | PLpro(8.75);nsp15(6.71) |
| K279-1256 | K279-1256 | | 958963-23-0 | (=C2C=C3) | O4S | 450.52 | proteolytic subunit | proteolytic subunit inhibitor | PLpro(8.75);X Domain(10.57) |
| K405-3134 | K405-3134 | | 890889-69-7 | NCCC(=CC | C21H25N5 | 347.47 | Microtubule-associated protein tau | inhibitor | PLpro(8.72) |
| E216-4969 | E216-4969 | | 891882-89-6 | C=CC=CC | O4 | 354.41 | Microtubule-associated protein tau | inhibitor | PLpro(8.7) |
| C547-0142 | C547-0142 | | 901729-44-0 | C(C=1OCC | O4 | 380.45 | Nonstructural protein 1 | Nonstructural protein 1 inhibitor | PLpro(8.68) |
| C880-2612 | C880-2612 | | 906252-12-8 | (C=NN1CC | N4O2 | 484.99 | Cellular tumor antigen p53 | Cellular tumor antigen p53 inhibitor | PLpro(8.65) |
| E542-1696 | E542-1696 | | 892370-71-7 | C1=C/C=C( | O2 | 440.59 | DNA polymerase beta | DNA polymerase beta inhibitor | PLpro(8.63) |
| T7210 | Guanosine 5'-diphosphate | GDP | 146-91-8 | nc2c(=O)[n | O11P2 | 443.2 | Endogenous Metabolite | Iron Mobilizer, Preventing the Hepcidin- | PLpro(8.62) |
| T5330 | Fluralaner | 3 | 864731-61-3 | C(=O)NCC( | F6N3O3 | 556.29 | GABA Receptor | ectoparasiticide. It potently and | PLpro(8.61) |
| 8009-8507 | 8009-8507 | | 309283-33-8 | )C=C(C(F)( | NOS | 435.47 | Androgen Receptor | Androgen Receptor inhibitor | PLpro(8.57) |
| T7509 | PD 117519 | CI947 | 96392-15-3 | 2cnc3c(NC | O4 | 383.4 | Adenosine Receptor | receptor | PLpro(8.54) |
| TMO2681 | Xanthosine | riboside;9-Beta- | 146-80-5 | O[C@H]([C | O6 | 284.22 | Others | induced germination of Bacillus | PLpro(8.54) |
| 6623-1226 | 6623-1226 | | 350497-73-3 | =C/C(=CC | O2S | 419.51 | Cellular tumor antigen p53 | Cellular tumor antigen p53 inhibitor | PLpro(8.53) |
| T0148L | Pentahydrate | calcium salt | 6035-45-6 | N1)[nH]c(n | N7O·5H2 | 601.58 | Others | used in combination with other | RdRP(11.73) |
| T6676 | Sofosbuvir | 7977 | 1190307-88-0 | )(OC[C@H] | 3O9P | 529.45 | HCV Protease inhibitor | analog inhibitor of hepatitis C virus | RdRP(11.49) |
| T4060 | Acelarin | NUC-1031 | 840506-29-8 | P(=O)(Oc1 | N4O8P | 580.47 | DNA/RNA Synthesis inhibitor | enhancement and transformation of | RdRP(11.46) |
| C241-1670 | C241-1670 | | 912800-78-3 | NC(=CC2[ | N4O6S | 583.07 | Thyroid hormone receptor beta-1 | inhibitor | RdRP(11.29) |
| T3893 | Forsythoside B | | 81525-13-5 | C@@H]([C | 9 | 756.71 | NF-κB inhibitor | reduces the biological activity of serum | RdRP(11.24) |
| T7086 | TBTA | | 510758-28-8 | Cc2ccccc2) | 0 | 530.62 | Others | 2, 3-triazole groups. It complexes with, | RdRP(11.15) |
| T3780 | Oroxin B | Hypocretin-2 | 114482-86-9 | O[C@@H]( | 5 | 594.52 | Others | Oroxin B has antioxidant activity. | RdRP(10.9) |
| C700-0693 | C700-0693 | | 902482-58-0 | C=3C=C2N | O6S | 508.64 | Ataxin-2 | Ataxin-2 inhibitor | RdRP(10.85) |
| C880-0271 | C880-0271 | | 906237-73-8 | CC2)C3)(N | O2 | 486.62 | Ataxin-2 | Ataxin-2 inhibitor | RdRP(10.77) |
| T5345 | V9302 | V 9302;V-9302 | 1855871-76-9 | Oc2ccccc2 | O4 | 538.69 | Others | antagonist of transmembrane | RdRP(10.77) |
| T0672 | Pravastatin sodium | (sodium);CS-514 | 81131-70-6 | @H](C[C@ | O7 | 446.52 | HMG-CoA Reductase inhibitor | reductase inhibitor, inhibits sterol | RdRP(10.69);X Domain(10.4) |
| T3670 | Forsythoside A | Forsythiaside | 79916-77-1 | 1O[C@@H | 5 | 624.59 | Others inhibitor | anticomplementary, anti-inflammatory | RdRP(10.67) |
| T4255 | TM5275 sodium | salt | 1103926-82-4 | ]C(=O)c4cc | N3NaO5 | 543.98 | PAI-1 | activator inhibitor 1 (PAI-1). | RdRP(10.53) |
| 6747-0106 | 6747-0106 | | | C@@]([H]) | O4 | 340.38 | Alpha-galactosidase A | Alpha-galactosidase A inhibitor | RdRP(10.42);nsp15(6.78) |

| | | | | | | | | | |
|---|---|---|---|---|---|---|---|---|---|
| E946-0779 | E946-0779 | | 950308-80-2 | )C(=NC=2 | O3 | 436.56 | factor 2 | factor 2 inhibitor | RdRP(10.38) |
| D072-0556 | D072-0556 | | 862742-56-1 | =CC2)C=C | N3O3S | 479.99 | homolog 3 | homolog 3 inhibitor | RdRP(10.35) |
| 6286-0223 | 6286-0223 | | 615280-85-8 | C(C1=CC1) | O2 | 400.53 | Inositol monophosphatase 1 | Inositol monophosphatase 1 inhibitor | RdRP(10.34) |
| C448-1053 | C448-1053 | | 901863-37-4 | N1)C(C=C2 | O3 | 475.6 | HEK293 | HEK293 inhibitor | RdRP(10.32) |
| T5416 | T-5224 | | 530141-72-1 | c1cc(ccc1O | 8 | 517.53 | MMP | Fos/AP-1 inhibitor, which specifically | RdRP(10.29) |
| C636-2422 | C636-2422 | | 1037192-29-2 | C(=CC2)C | 3O3 | 449.53 | HEK293 | HEK293 inhibitor | RdRP(10.26);nsp15(7.06) |
| C336-0089 | C336-0089 | | 1037293-40-5 | =N[C@]1([ | N5O3S | 538.07 | Ferritin light chain | Ferritin light chain inhibitor | RdRP(10.23) |
| T1938 | FLT3-IN-2 | 0 | 923562-23-6 | ccc(CNc2n | 3N4 | 416.83 | FLT inhibitor | μM). | RdRP(10.22) |
| T2132 | Buspirone hydrochloride | HCl;Buspar;Naro | 33386-08-2 | C2(CCCC2 | N5O2 | 421.96 | Receptor antagonist | agonist, used to treat generalized | RdRP(10.22) |
| T5847 | Cloprostenol sodium | sodium;ICI | 55028-72-3 | COc1cccc( | NaO6 | 446.9 | Prostaglandin Receptor | solute, crystalline form of cloprostenol | RdRP(10.21);X Domain(11.32) |
| T3263 | Cefminox Sodium | na;Tencef | 92636-39-0 | @]1(NC(= | Na2O7S3 | 563.53 | Antibiotic inhibitor | bactericidal cephalosporin antibiotic. It | RdRP(10.2) |
| 8539-0868 | 8539-0868 | | 950264-69-4 | )(N=C(C(C( | N3O4 | 463.92 | homolog 3 | homolog 3 inhibitor | RdRP(10.18) |
| T5384 | RS 504393 | | 300816-15-3 | CCN1CCC | O3 | 417.51 | CCR | chemokine receptor antagonist (IC50s: | RdRP(10.17) |
| K935-0047 | K935-0047 | | | ]([H])(C/C1 | N2O4 | 517.03 | Prelamin-A/C | Prelamin-A/C inhibitor | RdRP(10.05) |
| 8014-1195 | 8014-1195 | | 1008072-65-8 | (NC(C(=O) | O3 | 379.42 | Cellular tumor antigen p53 | Cellular tumor antigen p53 inhibitor | RdRP(10.04) |
| T2375 | BX471 | 471;ZK-811752 | 217645-70-0 | N(C(=O)C | N4O3 | 434.89 | CCR inhibitor | peptide CCR1 antagonist. | RdRP(10.04) |
| E843-0272 | E843-0272 | | 894932-72-0 | =CC2)C=C | O5 | 454.45 | proteolytic subunit | proteolytic subunit inhibitor | RdRP(10.03) |
| G768-1619 | G768-1619 | | 959516-20-2 | N=C1[S]C( | O3S | 504.62 | proteolytic subunit | proteolytic subunit inhibitor | RdRP(10.03) |
| T6179 | Latamoxef sodium | 6059S;LY- | 64953-12-4 | .CO[C@]1( | Na2O9S | 564.44 | Antibacterial inhibitor | compound more effective against | RdRP(10) |
| T5725 | lithospermic acid B | B;Salvianolic | 121521-90-2 | @@H](Cc1 | 6 | 718.61 | Sirtuin | antioxidant from Salvia extract. It plays | X Domain(14.2) |
| T0136 | Citicoline | Choline;cytidine | 987-78-0 | )CCOP(=O | O11P2 | 488.32 | Dopamine Receptor antagonist | synthesis of phosphatidylcholine, a | X Domain(12.27) |
| 8015-9350 | 8015-9350 | | 695204-55-8 | C=C(C(O2) | N3O5 | 457.92 | methyltransferase MLL | methyltransferase MLL inhibitor | X Domain(11.91) |
| 8015-7947 | 8015-7947 | | 696653-08-4 | C=C(C(O2) | O7 | 469.5 | factor 2 | factor 2 inhibitor | X Domain(11.69) |
| T0154 | Nebivolol hydrochloride | hydrochloride;Ne | 152520-56-4 | H](CNC[C | NO4·HCl | 441.9 | Adrenergic Receptor antagonist | cardioselective ADRENERGIC BETA-1 | X Domain(11.38);nsp15(7.55) |
| T3409 | Plantamajoside | Y0160;C10485 | 104777-68-6 | (C=C1CCO | 6 | 640.6 | Others | anti-inflammatory, antinociceptive | X Domain(11.21) |
| C528-0901 | C528-0901 | | 901875-64-7 | C=C2OC)= | O6S | 437.52 | Geminin | Geminin inhibitor | X Domain(11.16) |
| K284-3774 | K284-3774 | | 422285-64-1 | CC(=O)NC | 3O3S | 489.57 | Beta-lactamase AmpC | Beta-lactamase AmpC inhibitor | X Domain(11.11) |
| C527-0061 | C527-0061 | | 958954-53-5 | 1N=C(OC= | O6S | 442.49 | Beta-lactamase AmpC | Beta-lactamase AmpC inhibitor | X Domain(11.01) |
| G678-0299 | G678-0299 | | 904828-70-2 | CN1C(=CC | 3O5S | 465.55 | Geminin | Geminin inhibitor | X Domain(10.97) |
| C060-0100 | C060-0100 | | 443331-83-7 | (N=CN1CC | O5S | 525.63 | Beta-lactamase AmpC | Beta-lactamase AmpC inhibitor | X Domain(10.88) |
| T3708 | BP-1-102 | | 1334493-07-0 | N(Cc1ccc(c | N2O6S | 626.15 | STAT inhibitor | and specific STAT3 inhibitor. BP-1-102 | X Domain(10.84) |
| 7582-0307 | 7582-0307 | | 824980-59-8 | N(CC(=O)N | O3S2 | 407.56 | Beta-lactamase AmpC | Beta-lactamase AmpC inhibitor | X Domain(10.83) |
| C336-0168 | C336-0168 | | 959484-31-2 | (=C2C=C3) | 5O5S | 559.58 | homolog 3 | homolog 3 inhibitor | X Domain(10.76);nsp15(6.88) |
| T3671 | Vitexin-2''-O-rhamnoside | rhaMnoside;2''-O- | 64820-99-1 | 1O[C@@H | 4 | 578.52 | Others | compound contributes to the protection | X Domain(10.53) |
| G205-0745 | G205-0745 | | 951489-25-1 | 2)(N=C1N( | O4S | 443.53 | proteolytic subunit | proteolytic subunit inhibitor | X Domain(10.5) |
| Y041-2269 | Y041-2269 | | | =CC(OCC( | O7 | 478.51 | Geminin | Geminin inhibitor | X Domain(10.49) |
| 7680-2228 | 7680-2228 | | 838252-73-6 | N(CC(=O)N | O5S2 | 466.58 | Histone acetyltransferase GCN5 | inhibitor | X Domain(10.37) |
| C200-1101 | C200-1101 | | 1037289-06-7 | =N[C@@]1 | N4O2S | 517.05 | FK506 binding protein 12 | FK506 binding protein 12 inhibitor | X Domain(10.37);nsp15(7.22) |
| T7142 | Cephalosporin C zinc salt | | 59143-60-1 | C@@]12S | O8SZn | 478.78 | Others | inhibitor of SAMHD1 (IC50 : 1.1 ± 0.1 | X Domain(10.36) |
| G856-4196 | G856-4196 | | | =C/C=C(/C | O4 | 486.58 | Prelamin-A/C | Prelamin-A/C inhibitor | X Domain(10.29) |
| T2402 | Tianeptine sodium | sodium salt | 30123-17-2 | c2cC(c2c(S1 | N2NaO4S | 458.93 | 5-HT Receptor agonist | serotonin reuptake enhancer (SSRE), | X Domain(10.26) |
| C906-0334 | C906-0334 | | 890800-06-3 | C2)CC2)(C | O3 | 423.56 | Plasmodium falciparum | Plasmodium falciparum inhibitor | X Domain(10.18);nsp15(6.84) |
| T2923 | Apremilast | CC-10004 | 608141-41-9 | c(c1)[C@ | O7S | 460.5 | PDE inhibitor | orally active PDE4 (IC50=74 nM) with | X Domain(10.09) |
| Y050-1938 | Y050-1938 | | 878423-88-2 | N(C)CC(=O | O6S | 406.46 | Ataxin-2 | Ataxin-2 inhibitor | X Domain(10.09) |
| T3879 | Silychristin | Silicristin | 33889-69-9 | COc1cc(cc | C25H22O1 | 482.44 | Lipoxygenase inhibitor | Silychristin is a plant growth regulator. | X Domain(10.05) |
| T1227 | Cepazine | axetil;Ceftin;Zinn | 64544-07-6 | (=O)N[C@ | O10S | 510.47 | Antibiotic | cephalosporin antibiotic. | X Domain(10.03) |
| K786-6600 | K786-6600 | | 697781-60-5 | =CC2)C=C | O2 | 406.53 | Microtubule-associated protein tau | inhibitor | X Domain(9.99) |
| E587-0421 | E587-0421 | | 878055-77-7 | (=C1)SCC( | O3S | 465.62 | homolog 3 | homolog 3 inhibitor | X Domain(9.98) |
| C651-0859 | C651-0859 | | 902582-18-7 | C2C=C(C( | O6 | 386.41 | Lysosomal alpha-glucosidase | Lysosomal alpha-glucosidase inhibitor | X Domain(9.97) |
| T1066 | Ketanserin | inum;Ketanserin | 74050-98-9 | )C(=O)C1C | 3O3 | 395.43 | 5-HT Receptor antagonist | and serotonin (5-hydroxytryptamine, | X Domain(9.92) |
| T1331 | Flavin mononucleotide | phosphate | 130-40-5 | nc3c(n1C[C | NaO9P | 478.33 | Vitamin inhibitor | soluble, essential micronutrient that is | X Domain(9.92) |
| E465-0564 | E465-0564 | | 891924-26-8 | CC1C(=O) | O5S2 | 398.5 | Beta-lactamase AmpC | Beta-lactamase AmpC inhibitor | X Domain(9.91) |
| T7100 | PLX-5622 | | 1303420-67-8 | )cc1CNc1c | N5O | 395.41 | CSF-1R | penetrant and oral active CSF1R | X Domain(9.9) |
| E986-1019 | E986-1019 | | 894894-84-9 | =O)NC/C1 | O6 | 477.52 | factor 2 | factor 2 inhibitor | X Domain(9.87) |
| C700-1423 | C700-1423 | | 902589-58-6 | CC=1N=C( | N3O5S | 441.94 | Histone acetyltransferase GCN5 | inhibitor | X Domain(9.77) |
| C636-1184 | C636-1184 | | 1037192-50-9 | =O)[C@@] | O5 | 477.57 | Cellular tumor antigen p53 | Cellular tumor antigen p53 inhibitor | X Domain(9.74) |
| E461-0614 | E461-0614 | | 872840-60-3 | C1(=C(N(C | C20H23N5 | 397.44 | Cellular tumor antigen p53 | Cellular tumor antigen p53 inhibitor | nsp15(8.58) |
| E461-0573 | E461-0573 | | 300394-36-9 | C1(=C(N(C | C19H21N5 | 383.41 | Beta-lactamase AmpC | Beta-lactamase AmpC inhibitor | nsp15(8.25) |
| T5171 | Treprostinil Sodium | UT-15 | 289480-64-4 | [Na+].CCC | C23H33Na | 412.5 | Prostaglandin Receptor;VEGFR;c-RET | Treprostinil is a potent DP1, IP and | nsp15(8.17) |
| T1503 | Esmolol hydrochloride | Esmolol | 81161-17-3 | c1(ccc(cc1) | C16H25NO | 331.15 | Adrenergic Receptor antagonist | Esmolol is a cardioselective beta- | nsp15(7.98) |
| T3886 | Rosavin | Rosavidin | 84954-92-7 | O1[C@H]([ | C20H28O1 | 428.43 | P450 inhibitor | Rosavin has antidepressant and | nsp15(7.94) |
| G948-4026 | G948-4026 | | 933019-97-7 | O=[S](=O)( | C24H28N4 | 468.58 | Ferritin light chain | Ferritin light chain inhibitor | nsp15(7.83) |
| T2911 | Stevioside | | 57817-89-7 | C[C@@]12 | C38H60O1 | 804.88 | TLR inhibitor | A natural, noncaloric sweetener with a | nsp15(7.72) |
| C700-1219 | C700-1219 | | 902443-53-2 | S(C(C=C(N | C28H36N4 | 540.69 | Mothers against decapentaplegic | Mothers against decapentaplegic | nsp15(7.7) |
| Y031-0649 | Y031-0649 | | 725693-58-3 | S(NCC(OC | C18H19FN | 378.43 | Geminin | Geminin inhibitor | nsp15(7.68) |
| T6085 | PF543 | PF 543;PF- | 1415562-82-1 | Cc1cc(CS( | C27H31NO | 465.6 | S1P Receptor inhibitor | PF-543, a novel sphingosine- | nsp15(7.66) |
| Y031-0429 | Y031-0429 | | 725693-41-4 | S(NCC(OC | C18H19FN | 378.43 | Nuclear receptor ROR-gamma | Nuclear receptor ROR-gamma inhibitor | nsp15(7.66) |
| T3899 | Calceolarioside B | Nuomioside | 105471-98-5 | C1=CC(=C | C23H26O1 | 478.45 | Others | Calceolarioside B displays inhibition of | nsp15(7.63) |
| C880-0694 | C880-0694 | | 906243-33-2 | O=C1C4=C | C26H27N5 | 457.54 | Guanine nucleotide-binding protein | Guanine nucleotide-binding protein | nsp15(7.59) |
| T6846 | Vesatolimod | GS-9620 | 1228585-88-3 | CCCCOc1n | C22H30N6 | 410.51 | TLR inhibitor | GS-9620 is an effective and specific | nsp15(7.55) |
| D715-0611 | D715-0611 | | 956040-89-4 | C(O1)C(C( | C25H25NO | 435.48 | Plasmodium falciparum | Plasmodium falciparum inhibitor | nsp15(7.54) |
| D134-0324 | D134-0324 | | 876725-25-6 | O=C(NCC= | C18H19N3 | 341.37 | Histone-lysine N-methyltransferase, H3 | Histone-lysine N-methyltransferase, H3 | nsp15(7.5) |
| Y031-2258 | Y031-2258 | | 838871-13-9 | N1C=C(C( | C19H19Br | 387.28 | Nonstructural protein 1 | Nonstructural protein 1 inhibitor | nsp15(7.49) |
| T7413 | JNJ-5207852 | | 398473-34-2 | C(COc1ccc | C20H32N2 | 316.48 | Histamine Receptor | JNJ-5207852 is a novel, non-imidazole | nsp15(7.48) |
| T0342 | Carvedilol phosphate | BM 14190 | 610309-89-2 | O.OP(O)(O | C24H26N2 | 522.49 | Adrenergic Receptor antagonist;Gap | Carvedilol Phosphate is the phosphate | nsp15(7.48) |
| D389-0267 | D389-0267 | | 848205-63-0 | C1(=C(N(C | C26H27N5 | 473.54 | Nonstructural protein 1 | Nonstructural protein 1 inhibitor | nsp15(7.47) |
| 6913-0019 | 6913-0019 | | 909860-72-6 | C(=C(N1C) | C22H32N2 | 388.51 | Beta-lactamase AmpC | Beta-lactamase AmpC inhibitor | nsp15(7.46) |
| T6S0119 | Dauricine | | 524-17-4 | COc1cc2C | C38H44N2 | 624.77 | Others | 1. Dauricine has pulmonary toxicity, | nsp15(7.44) |
| T7183 | CPI-444 | V81444;ciforade | 1202402-40-1 | Cc1ccc(o1)- | C20H21N7 | 407.43 | Adenosine Receptor | CPI-444 is an antagonist of the | nsp15(7.44) |
| G008-5517 | G008-5517 | | 894918-95-7 | S(C(=CC1 | C26H33N3 | 499.63 | Glucagon-like peptide 1 receptor | Glucagon-like peptide 1 receptor | nsp15(7.42) |
| T0179 | Ticagrelor | AZD6140;AR-C | 274693-27-5 | CCCSc1nc | C23H28F2 | 522.57 | Others antagonist; P2 Receptor | Ticagrelor, produced by AstraZeneca, | nsp15(7.42) |
| E977-0894 | E977-0894 | | 894887-97-9 | O=[S](=O)( | C21H25N3 | 431.51 | Thyroid hormone receptor beta-1 | Thyroid hormone receptor beta-1 | nsp15(7.42) |
| 6655-0470 | 6655-0470 | | 760196-67-6 | O=C(/C2= | C22H19NO | 361.47 | Inositol monophosphatase 1 | Inositol monophosphatase 1 inhibitor | nsp15(7.4) |
| T4S0998 | Trifolirhizin | | 6807-83-6 | OC[C@H]6 | C22H22O1 | 446.4 | Adrenergic Receptor;TNF | Trifolirhizin exerts varying degrees of | nsp15(7.34) |
| T5S1094 | Forsythoside E | | 93675-88-8 | C[C@@H] | C20H30O1 | 462.44 | Others | Forsythoside E is a natutal product | nsp15(7.33) |
| T4602 | Hydrocortisone 21- | Hydrocortisone | 2203-97-6 | [H][C@@]1 | C25H34O8 | 462.53 | Others | Hydrocortisone 21-hemisuccinate is a | nsp15(7.32) |
| E977-0921 | E977-0921 | | 894888-70-1 | O=[S](=O)( | C21H25N3 | 415.52 | Nuclear factor erythroid 2-related | Nuclear factor erythroid 2-related | nsp15(7.32) |
| G937-2830 | G937-2830 | | 933202-35-8 | O=[S](=O)( | C22H29Cl | 497.02 | Prelamin-A/C | Prelamin-A/C inhibitor | nsp15(7.31) |
| F070-0397 | F070-0397 | | 959506-77-5 | C(C(=N1)N | C25H27F3 | 488.51 | Cellular tumor antigen p53 | Cellular tumor antigen p53 inhibitor | nsp15(7.29) |
| T0812 | Propranolol hydrochloride | Propranolol | 318-98-9 | Cl.CC(C)N | C16H22Cl | 295.81 | Adrenergic Receptor antagonist | Propranolol hydrochloride is a widely | nsp15(7.27) |
| G357-1938 | G357-1938 | | 959562-33-5 | C(N(C1=O) | C24H21N3 | 463.52 | Glucagon-like peptide 1 receptor | Glucagon-like peptide 1 receptor | nsp15(7.27) |
| 5228-0298 | 5228-0298 | | 476481-85-3 | C(=C(C1= | C19H24N4 | 388.49 | Plasmodium falciparum | Plasmodium falciparum inhibitor | nsp15(7.25) |
| E545-0290 | E545-0290 | | 892427-14-4 | O=C3N(CC | C22H22N2 | 362.43 | Nuclear factor erythroid 2-related | Nuclear factor erythroid 2-related | nsp15(7.25) |
| T1959 | BIX 01294 Trihydrochloride | BIX 01294 | 1392399-03-9 | Cl.COc1c( | C28H38N6 | 600.02 | Histone Methyltransferase inhibitor | BIX01294 is an inhibitor of G9a histone | nsp15(7.25) |
| C199-0140 | C199-0140 | | 850935-38-5 | N(C(C(C(= | C28H33NO | 479.58 | Mothers against decapentaplegic | Mothers against decapentaplegic | nsp15(7.24) |
| N081-0783 | N081-0783 | | 332849-41-9 | C(C(CC(=O | C20H21NO | 355.39 | Beta-lactamase AmpC | Beta-lactamase AmpC inhibitor | nsp15(7.22) |
| C258-0605 | C258-0605 | | 862316-96-9 | C(N(C1=O) | C28H29N3 | 487.63 | TAR DNA-binding protein 43 | TAR DNA-binding protein 43 inhibitor | nsp15(7.21) |
| C199-0115 | C199-0115 | | 850935-47-6 | N(C(C(=CC | C30H29NO | 499.57 | Ataxin-2 | Ataxin-2 inhibitor | nsp15(7.2) |

| | | | | | | | | | |
|---|---|---|---|---|---|---|---|---|---|
| D011-0852 | D011-0852 | | 883637-27-2 | O=C1C[C | C27H26Cl | 475.98 | Cellular tumor antigen p53 | Cellular tumor antigen p53 inhibitor | nsp15(7.2) |
| 1349-0007 | 1349-0007 | | 314033-58-4 | O=C(O)CC | C18H18N2 | 342.42 | ATPase family AAA domain-containing | ATPase family AAA domain-containing | nsp15(7.2) |
| 5782-5442 | 5782-5442 | | 496774-72-2 | C(=CC1C( | C23H17NO | 403.46 | Microtubule-associated protein tau | Microtubule-associated protein tau | nsp15(7.16) |
| C528-1116 | C528-1116 | | 901742-07-2 | C(=NC(CS( | C21H28N2 | 404.53 | Guanine nucleotide-binding protein | Guanine nucleotide-binding protein | nsp15(7.16) |
| K405-3034 | K405-3034 | | 890884-23-8 | N(C(=C1C2 | C22H23N5 | 357.46 | Microtubule-associated protein tau | Microtubule-associated protein tau | nsp15(7.14) |
| T5S0733 | Picroside III | | 64461-95-6 | COc1cc(\C | C25H30O1 | 538.5 | Others | Picroside III, an iridoid glucoside found | nsp15(7.14) |
| T6018 | Zosuquidar 3HCl | LY-335979 | 167465-36-3 | Cl.Cl.Cl.O[ | C32H31F2 | 636.99 | P-gp modulator | Zosuquidar (LY335979) is a potent | nsp15(7.14) |
| T2018 | Apilimod | STA 5326 | 541550-19-0 | c1(nc(cc(n1 | C23H26N6 | 418.49 | IL Receptor inhibitor;PI3K | Apilimod inhibits the production of IL- | nsp15(7.13) |
| G361-0500 | G361-0500 | | 932281-65-7 | C(N(N1)C2 | C24H32N6 | 436.56 | Glucagon-like peptide 1 receptor | Glucagon-like peptide 1 receptor | nsp15(7.13) |
| C450-0646 | C450-0646 | | 1034591-56-4 | S(NC(C(NC | C24H31N3 | 473.6 | Nuclear factor erythroid 2-related | Nuclear factor erythroid 2-related | nsp15(7.12) |
| K786-2099 | K786-2099 | | 1037168-03-8 | O=C2N(C/ | C28H30N2 | 458.56 | Cellular tumor antigen p53 | Cellular tumor antigen p53 inhibitor | nsp15(7.11) |
| 8388-0819 | 8388-0819 | | | O=C(C[S]C | C15H18Cl | 376.31 | Microtubule-associated protein tau | Microtubule-associated protein tau | nsp15(7.1) |
| 3257-2542 | 3257-2542 | | 292612-55-6 | O=C(OCC( | C25H18Cl | 415.88 | Microtubule-associated protein tau | Microtubule-associated protein tau | nsp15(7.1) |
| D236-0021 | D236-0021 | | 1008958-65-3 | C(C(CC(=O | C15H20N2 | 276.34 | Guanine nucleotide-binding protein | Guanine nucleotide-binding protein | nsp15(7.07) |
| T0228 | Methyl hesperidin | | 11013-97-1 | COc1ccc(c | C29H36O1 | 624.59 | Akt inhibitor; PKC inhibitor | Methyl Hesperidin, a flavanone | nsp15(7.06) |
| E750-0072 | E750-0072 | | 1037256-79-3 | N(C(N1C(C | C23H24FN | 409.46 | Mothers against decapentaplegic | Mothers against decapentaplegic | nsp15(7.06) |
| K279-0833 | K279-0833 | | 1037168-66-3 | O=C1N4C( | C29H28N4 | 512.64 | Prelamin-A/C | Prelamin-A/C inhibitor | nsp15(7.05) |
| T6633 | Ranolazine | RS 43285- | 95635-55-5 | COc1ccccc | C24H33N3 | 427.54 | Calcium Channel inhibitor | Ranolazine is a calcium uptake | nsp15(7.04) |
| T5711 | Methylnissolin-3-O- | (6aR, 11 aR)-3- | 94367-42-7 | [H][C@@]1 | C23H26O1 | 462.45 | Others | Methylnissolin-3-O-glucoside has anti- | nsp15(7.02) |
| D389-0835 | D389-0835 | | 877803-29-7 | C(=C(C1= | C21H25N5 | 411.46 | HEK293 | HEK293 inhibitor | nsp15(7.01) |
| 8009-9265 | 8009-9265 | | 309278-11-3 | O=[S](=O)( | C22H22N2 | 410.5 | Cannabinoid CB1 receptor | Cannabinoid CB1 receptor inhibitor | nsp15(7.01) |
| C795-0720 | C795-0720 | | 903150-68-5 | N(C(C(N(N | C20H29N5 | 403.55 | Aberrant vpr protein | Aberrant vpr protein inhibitor | nsp15(7.01) |
| C320-0115 | C320-0115 | | 688759-32-2 | N(C(=N1)C | C21H31N5 | 401.51 | Plasmodium falciparum | Plasmodium falciparum inhibitor | nsp15(7.01) |
| T3540 | IMR-1A | | 331862-41-0 | COc1cc(\C | C13H11NO | 325.36 | Gamma-secretase inhibitor | IMR-1A is the metabolite of IMR-1 | nsp15(7) |
| E544-0037 | E544-0037 | | 892379-45-2 | C(=C(C1S | C27H21F3 | 538.55 | Mothers against decapentaplegic | Mothers against decapentaplegic | nsp15(6.99) |
| T2306 | Brexpiprazole | OPC-34712 | 913611-97-9 | O=c1ccc2c | C25H27N3 | 433.57 | 5-HT Receptor agonist; Adrenergic | Brexpiprazole is a partial agonist of | nsp15(6.98) |
| C796-1295 | C796-1295 | | 959515-28-7 | O=C(C[S]/ | C26H24ClF | 495.02 | Survival motor neuron protein | Survival motor neuron protein inhibitor | nsp15(6.98) |
| T2491 | AZ5104 | | 1421373-98-9 | CN(C)CCN | C27H31N7 | 485.58 | EGFR inhibitor | AZ5104 is a potent EGFR inhibitor. | nsp15(6.98) |
| D364-2068 | D364-2068 | | 1147183-28-5 | N(C(N1)=O | C27H29N3 | 491.61 | TAR DNA-binding protein 43 | TAR DNA-binding protein 43 inhibitor | nsp15(6.97) |
| T3S1149 | Ganoderic acid G | | 98665-22-6 | C[C@H](C | C30H44O8 | 532.67 | Others | Ganoderic acid G is a highly oxidized | nsp15(6.96) |
| C848-0215 | C848-0215 | | 863449-34-7 | C(N=C1S(= | C18H24N2 | 428.59 | Glucagon-like peptide 1 receptor | Glucagon-like peptide 1 receptor | nsp15(6.95) |
| T0696 | Naftopidil | BM-15275;KT- | 57149-07-2 | COc1ccccc | C24H28N2 | 392.49 | Adrenergic Receptor antagonist | Naftopidil (INN, marketed under the | nsp15(6.94) |
| G856-5036 | G856-5036 | | 877642-30-3 | O=[S](=O)( | C21H20N4 | 472.55 | Cellular tumor antigen p53 | Cellular tumor antigen p53 inhibitor | nsp15(6.94) |
| 8211-0295 | 8211-0295 | | 892720-69-3 | C(=C(N1)C | C21H22Cl | 367.88 | Glucagon-like peptide 1 receptor | Glucagon-like peptide 1 receptor | nsp15(6.92) |
| T6078 | Saracatinib | AZD0530 | 379231-04-6 | CN1CCN(C | C27H32Cl | 542.03 | BTK; c-Kit; EGFR; Src | Saracatinib (AZD0530) is an effective | nsp15(6.91) |
| 5678-0009 | 5678-0009 | | 685122-34-3 | C(N=C1C( | C22H16ClF | 442.9 | Bromodomain adjacent to zinc finger | Bromodomain adjacent to zinc finger | nsp15(6.9) |
| G408-1998 | G408-1998 | | 932331-05-0 | S(NC(C=C | C20H18N2 | 382.44 | Cellular tumor antigen p53 | Cellular tumor antigen p53 inhibitor | nsp15(6.9) |
| T2320 | Indacaterol | | 312753-06-3 | CCc1c(cc2 | C24H28N2 | 392.49 | Adrenergic Receptor agonist | Indacaterol (Onbrez; Arcapta) is a β2- | nsp15(6.88) |
| 4341-0035 | 4341-0035 | | 431934-42-8 | O=C2N(CC | C26H30N2 | 466.54 | Nuclear receptor ROR-gamma | Nuclear receptor ROR-gamma inhibitor | nsp15(6.88) |
| G071-0411 | G071-0411 | | 895100-88-6 | O=[S]3(=O) | C26H20F2 | 538.6 | Nuclear factor erythroid 2-related | Nuclear factor erythroid 2-related | nsp15(6.87) |
| T4302 | iCRT3 | | 901751-47-1 | O=C(NCC | C23H26N2 | 394.53 | Wnt/beta-catenin | iCRT3 is a Wnt and β-catenin- | nsp15(6.86) |
| T6091 | CP673451 | CP 673451;CP- | 343787-29-1 | N1(CCC(C | C24H27N5 | 417.5 | c-Kit inhibitor; PDGFR inhibitor; | CP-673451 is a specific inhibitor of | nsp15(6.86) |
| 6872-0294 | 6872-0294 | | 847159-57-3 | O=C3C=2 | C25H26N2 | 434.5 | Geminin | Geminin inhibitor | nsp15(6.85) |
| G361-0499 | G361-0499 | | 932335-20-1 | C(N(N1)C2 | C22H28N6 | 408.51 | Beta-lactamase AmpC | Beta-lactamase AmpC inhibitor | nsp15(6.84) |
| E570-2192 | E570-2192 | | 892751-50-7 | S(C1=CC2 | C21H31N3 | 421.56 | Plasmodium falciparum | Plasmodium falciparum inhibitor | nsp15(6.84) |
| K788-6635 | K788-6635 | | 899198-29-9 | O=C1CCC( | C25H31N3 | 437.54 | Prelamin-A/C | Prelamin-A/C inhibitor | nsp15(6.84) |
| T4050 | GSK2981278 | ROR gama | 1474110-21-8 | OCc1cc(S( | C25H35NO | 461.61 | ROR agonist | GSK2981278 is a highly potent and | nsp15(6.83) |
| D345-0032 | D345-0032 | | 1207659-19-5 | N(C(COC(= | C20H25N3 | 355.44 | FK506 binding protein 12 | FK506 binding protein 12 inhibitor | nsp15(6.82) |
| D072-1432 | D072-1432 | | 380190-61-4 | O=[S](=O)( | C23H26Cl | 476 | Histone acetyltransferase GCN5 | Histone acetyltransferase GCN5 | nsp15(6.82) |
| 2191-2709 | 2191-2709 | | 312718-81-3 | O=C2NC(C | C22H24N2 | 412.45 | Geminin | Geminin inhibitor | nsp15(6.81) |
| T3274 | S49076 | | 1265965-22-7 | O=C1N(Cc | C22H22N4 | 438.13 | c-Met/HGFR inhibitor; FGFR inhibitor; | S49076 is a novel, potent inhibitor of | nsp15(6.81) |
| C276-0156 | C276-0156 | | 725687-57-0 | C(C(=C1C( | C21H25NO | 371.44 | Nonstructural protein 1 | Nonstructural protein 1 inhibitor | nsp15(6.81) |
| C891-1711 | C891-1711 | | 890786-93-3 | O=C1C=4C | C23H28FN | 441.51 | Ataxin-2 | Ataxin-2 inhibitor | nsp15(6.81) |
| G678-0075 | G678-0075 | | 904826-87-5 | S(N(CC1)C | C22H28Cl | 482 | Prelamin-A/C | Prelamin-A/C inhibitor | nsp15(6.8) |
| K784-6292 | K784-6292 | | 1028072-39-0 | N(C(C1)=O | C25H29FN | 424.52 | Plasmodium falciparum | Plasmodium falciparum inhibitor | nsp15(6.79) |
| 8104-06075 | 8104-06075 | | 880808-69-5 | CN1N=N/N | C20H25N5 | 435.98 | FK506 binding protein 12 | FK506 binding protein 12 inhibitor | nsp15(6.78) |
| T1533 | Valganciclovir hydrochloride | Valganciclovir | 175865-59-5 | Cl.CC(C)[C | C14H23Cl | 390.82 | Others | Valganciclovir Hydrochloride is a | nsp15(6.77) |
| K786-7206 | K786-7206 | | 697781-61-6 | O=[S](=O)( | C23H35N3 | 497.62 | Beta-lactamase AmpC | Beta-lactamase AmpC inhibitor | nsp15(6.77) |
| G503-0141 | G503-0141 | | 932502-72-2 | O=[S](=O)( | C22H22N2 | 426.56 | Prelamin-A/C | Prelamin-A/C inhibitor | nsp15(6.76) |
| C651-0863 | C651-0863 | | 902289-38-7 | C(C(N1C(C | C23H24N4 | 436.47 | DNA polymerase iota | DNA polymerase iota inhibitor | nsp15(6.76) |
| 1630-1629 | 1630-1629 | | | O=C2CC(C | C24H32N2 | 412.53 | Thyroid stimulating hormone receptor | Thyroid stimulating hormone receptor | nsp15(6.75) |
| T2795 | Amygdalin | Laetrile | 29883-15-6 | c1ccc(cc1)[ | C20H27NO | 457.43 | Others | Amygdalin has antifibrotic, antitumor, | nsp15(6.75) |
| K279-1018 | K279-1018 | | 1037222-80-2 | N(C(=N1)C | C30H35N5 | 545.71 | HEK293 | HEK293 inhibitor | nsp15(6.74) |
| T0990 | Droperidol | Dehydrobenzperi | 548-73-2 | Fc1ccc(cc1 | C22H22FN | 379.43 | Dopamine Receptor antagonist | Droperidol is a Dopamine-2 Receptor | nsp15(6.74) |
| C919-0592 | C919-0592 | | 890832-00-5 | C(N(N1)C( | C26H31N5 | 445.57 | Cellular tumor antigen p53 | Cellular tumor antigen p53 inhibitor | nsp15(6.74) |
| T0216 | Eletriptan HBr | Eletriptan | 177834-92-3 | CN1CCC[C | C22H27Br | 463.43 | 5-HT Receptor agonist | Eletriptan hydrobromide is an orally | nsp15(6.73) |
| C328-0251 | C328-0251 | | 851860-70-3 | O(C(=N1)S | C18H21FN | 392.46 | Guanine nucleotide-binding protein | Guanine nucleotide-binding protein | nsp15(6.73) |
| T2899 | Liquiritin | Liquiritoside;Liqu | 551-15-5 | C1[C@H]( | C21H22O9 | 418.39 | Others | Liquiritin (LIQ) is a main component | nsp15(6.72) |
| 8011-7222 | 8011-7222 | | 380581-14-6 | C(C(O1)=O | C17H20N2 | 348.42 | TAR DNA-binding protein 43 | TAR DNA-binding protein 43 inhibitor | nsp15(6.72) |
| K786-6703 | K786-6703 | | 898216-05-2 | O=C2COC | C24H29N3 | 407.52 | Huntingtin | Huntingtin inhibitor | nsp15(6.71) |
| T6853 | GSK591 | GSK3203591;EP | 1616391-87-7 | O=C(NC[C | C22H28N4 | 380.48 | Histone Methyltransferase inhibitor | GSK591, Alternative Names are | nsp15(6.71) |
| K221-2578 | K221-2578 | | 896371-47-4 | O=C2NC=1 | C26H29Cl | 513 | Microtubule-associated protein tau | Microtubule-associated protein tau | nsp15(6.71) |
| D345-0054 | D345-0054 | | | O=C(N2CC | C21H26Cl | 424.37 | Prelamin-A/C | Prelamin-A/C inhibitor | nsp15(6.7) |
| K906-2514 | K906-2514 | | 899717-59-0 | C(=C1N(C | C30H36N4 | 500.65 | Neuropeptide S receptor | Neuropeptide S receptor inhibitor | nsp15(6.7) |
| T6S0444 | Salvianolic acid A | Dan Phenolic | 96574-01-5 | OC(=O)[C | C26H22O1 | 494.45 | MMP | Salvianolic acid A could protect the | nsp15(6.7) |
| K784-6600 | K784-6600 | | 1035375-06-4 | N(C(C(N(C | C26H34N2 | 454.57 | Beta-lactamase AmpC | Beta-lactamase AmpC inhibitor | nsp15(6.7) |
| G242-0650 | G242-0650 | | 932283-47-1 | O=[S](=O)( | C20H27N3 | 389.52 | Geminin | Geminin inhibitor | nsp15(6.69) |
| K279-1177 | K279-1177 | | 958612-96-9 | O=C1N4C( | C25H25Cl | 481.02 | Microtubule-associated protein tau | Microtubule-associated protein tau | nsp15(6.69) |
| E750-0832 | E750-0832 | | 1032119-53-1 | O=C2N(C( | C25H23N3 | 461.48 | Histone acetyltransferase GCN5 | Histone acetyltransferase GCN5 | nsp15(6.69) |
| G069-0064 | G069-0064 | | 959519-68-7 | O=[S](=O)( | C23H27N3 | 457.55 | Nuclear factor erythroid 2-related | Nuclear factor erythroid 2-related | nsp15(6.68) |
